# Single cell epigenomic atlas of the developing human brain and organoids

**DOI:** 10.1101/2019.12.30.891549

**Authors:** Ryan S. Ziffra, Chang N. Kim, Amy Wilfert, Tychele N. Turner, Maximilian Haeussler, Alex M. Casella, Pawel F. Przytycki, Anat Kreimer, Katherine S. Pollard, Seth A. Ament, Evan E. Eichler, Nadav Ahituv, Tomasz J. Nowakowski

## Abstract

Dynamic changes in chromatin accessibility coincide with important aspects of neuronal differentiation, such as fate specification and arealization and confer cell type-specific associations to neurodevelopmental disorders. However, studies of the epigenomic landscape of the developing human brain have yet to be performed at single-cell resolution. Here, we profiled chromatin accessibility of >75,000 cells from eight distinct areas of developing human forebrain using single cell ATAC-seq (scATACseq). We identified thousands of loci that undergo extensive cell type-specific changes in accessibility during corticogenesis. Chromatin state profiling also reveals novel distinctions between neural progenitor cells from different cortical areas not seen in transcriptomic profiles and suggests a role for retinoic acid signaling in cortical arealization. Comparison of the cell type-specific chromatin landscape of cerebral organoids to primary developing cortex found that organoids establish broad cell type-specific enhancer accessibility patterns similar to the developing cortex, but lack many putative regulatory elements identified in homologous primary cell types. Together, our results reveal the important contribution of chromatin state to the emerging patterns of cell type diversity and cell fate specification and provide a blueprint for evaluating the fidelity and robustness of cerebral organoids as a model for cortical development.

## Main text

The diverse cell types of the human cerebral cortex (Fig. 1a) have been mostly classified based on a handful of morphological, anatomical, and physiological features. Recent innovations in single cell genomics, such as single cell mRNA sequencing (scRNA-seq), have enabled massively parallel profiling of thousands of molecular features in every cell, uncovering the remarkable molecular diversity of cell types previously considered homologous, such as excitatory neurons located in different areas of the cerebral cortex^1–6^. However, the developmental mechanisms underlying the emergence of distinct cellular identities are largely unknown, as most cortical neurons are generated at stages that are inaccessible to experimentation^5^.

**Fig. 1:**
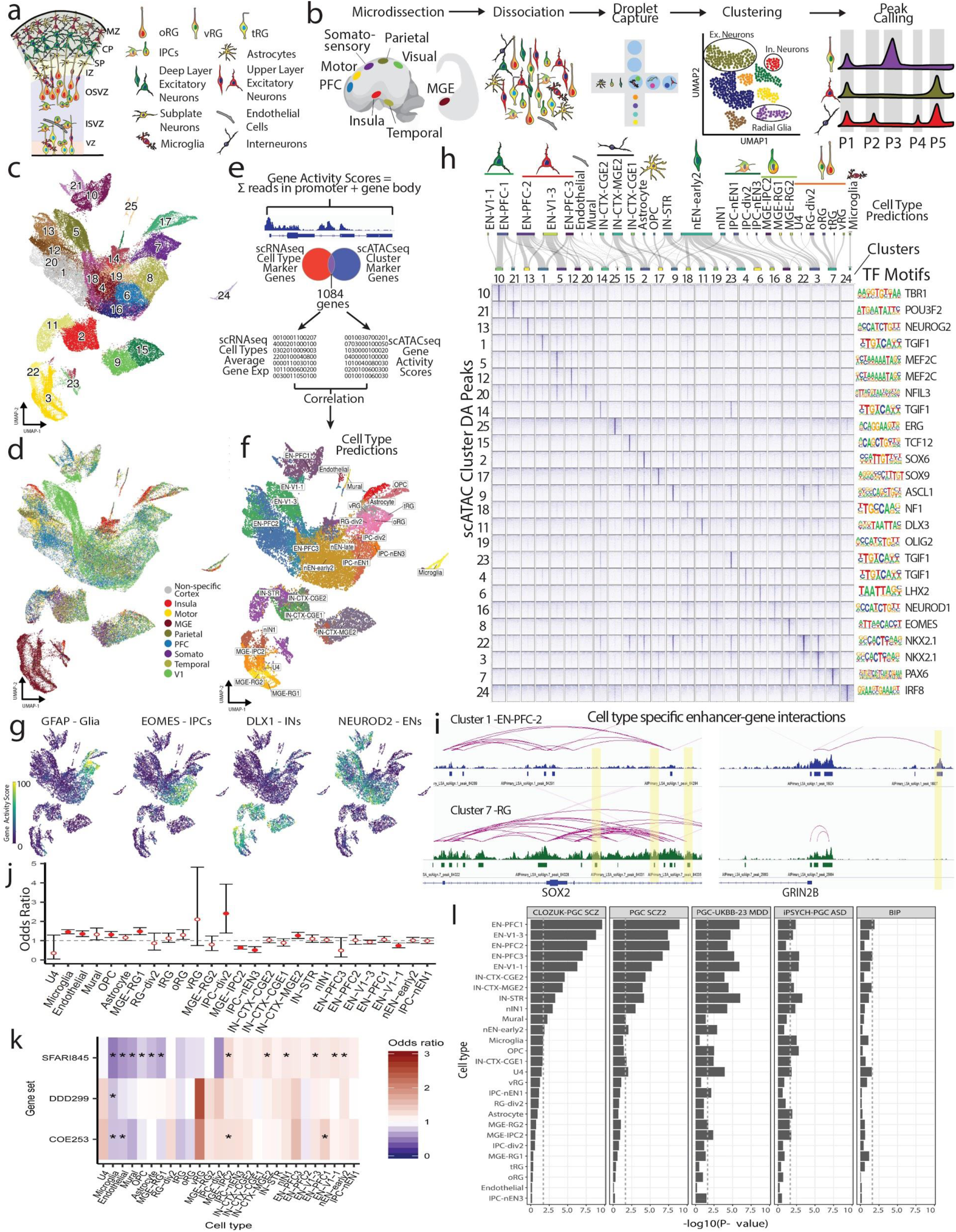
Single cell chromatin state atlas of the developing human brain. **a)** Schematic cross-section of developing cortex highlighting major cell types. VZ – ventricular zone, ISVZ – inner subventricular zone, OSVZ outer subventricular zone, IZ – intermediate zone, SP – subplate, CP-cortical plate, MZ – marginal zone, oRG outer radial glia, tRG – truncated radial glia, vRG – ventricular radial glia, IPCs – intermediate progenitor cells. **b)** Schematic depicting experimental workflow. PFC – prefrontal cortex, MGE – medial ganglionic eminence. **c)** UMAP projection of all primary scATAC-seq cells (n = 6 individuals, 77,354 cells) colored by leiden clusters. **d)** UMAP projection of all primary scATAC-seq cells colored by brain region of origin. Somato – somatosensory cortex, V1 – primary visual cortex. **e)** Gene activity scores were calculated for all protein coding genes for each cell based on summing reads in the promoter plus gene bodies. An intersecting set of 1084 marker genes was used to correlate gene activity scores to scRNA-seq data generated from single samples. Cell type predictions for scATACseq cells were made based on the max correlation values with cell type averages from the scRNAseq data (methods). **f)** UMAP projection of all primary scATAC-seq cells colored by cell type prediction. **g)** UMAP projections of gene activity scores for GFAP marking glia, EOMES marking intermediate progenitors, DLX1 marking cells in the interneuron lineage, and NEUROD2 marking cells in the excitatory neuron lineage. **h)** Top, sankey plot depicting mapping between scATACseq clusters and cell type predictions. Bottom left, Pileups of cluster specific ATAC-seq signal for each cluster within sets of the 1000 cluster specific peaks for each cluster by p-value (Fisher’s Exact, two-sided). Pileups are centered on peak centers and the +/-10Kb flanking region is depicted. Bottom right, significantly enriched TF motifs for each cluster specific peak set as determined by HOMER. **i)** Left, enhancer-gene interaction predictions for cluster 1 cells (EN-PFC-2) and cluster 7 cells (RG) in the SOX2 locus. Differentially accessible peaks that interact with SOX2 highlighted in yellow. Right, enhancer-gene interaction predictions for cluster 1 cells (EN-PFC-2) and cluster 7 cells (RG) in the GRIN2B locus, highlighting a peak interacting with GRIB2B in neurons but not in RGs. **j)** Enrichment and depletion of cell type-specific scATACseq peaks in copy number variant (CNV) regions enriched in pediatric cases of neurodevelopmental delay (NDD). Filled circles indicate Bonferroni corrected significance. **k)** Enrichment and depletion of scATACseq peaks in promoter and gene regions of genes associated for autism and NDD including genes enriched in *de novo* non-coding mutations (DNM) (SFARI845, DDD299^60^, COE253^61^). Stars indicate tests that pass Bonferroni significance. **l)** Heritability enrichment based on LD score regression analysis of GWAS summary statistics in cell type-specific peak sets from primary scATAC cells. From left to right, Pychiatric Genomics Consortium (PGC) schizophrenia (SCZ) GWAS^37^, an additional PGC schizophrenia GWAS^62^, PGC autism spectrum disorder (ASD) GWAS^63^, PGC major depressive disorder (MDD) GWAS^37^, PGC bipolar (BIP) disorder^64^.

Over 60 years ago, Conrad Waddington introduced the concept of an epigenomic landscape to account for the emergence of distinct cell fates^7^. In particular, chromatin state defines the functional architecture of the genome by modulating the accessibility of gene regulatory elements, such as enhancers, which serve as binding sites for transcriptional regulators. Together with the expression of unique combinations of transcription factors, chromatin state is believed to represent the *cis*-regulatory ‘vocabulary’ of gene expression, which is a fundamental determinant of cell identity^8, 9^. However, studies of chromatin state in the developing brain have been largely limited as established methods for discovering gene regulatory elements, such as the assay for transposase-accessible chromatin using sequencing^10^ (ATACseq) or chromatin immunoprecipitation followed by sequencing^11^ (ChIP-seq), lacked cellular resolution and mainly focused on studies that enrich for broad cell classes revealing changes in regional patterning and neuronal differentiation^8, 12–17^. Several methods have been recently developed to enable profiling of the epigenomic landscape at single cell resolution, such as scATAC-seq^18, 19^, revealing many cell type-specific patterns of enhancer activity in the developing and adult mouse brain, as well as the adult human brain^20–22^. However, it is particularly important to characterize gene regulatory elements in their native context of the developing human tissue, as growing evidence has shown that genetic variants associated with psychiatric disease reside in evolutionary accelerated sequences that are putative neurodevelopmental enhancers^23–26^.

### Chromatin states define the major cell types in the developing brain

To characterize the chromatin state landscape of the developing human brain at single cell resolution, we performed scATACseq on primary samples of the human forebrain at midgestation (n = 6 individuals). For a subset of samples, we preserved the anatomical region of origin information (Extended Data Table 1), including dorsolateral prefrontal cortex (PFC), primary visual cortex (V1), primary motor cortex (M1), primary somatosensory cortex, dorso-lateral parietal cortex, temporal cortex, insular cortex, and the medial ganglionic eminence (MGE), a major source of cortical interneurons^27, 28^ (Fig. 1b, Extended Data Table 1).

We generated data from 77,354 cells passing quality control criteria (Methods, Extended Data Fig. 1a-d). To reduce the dimensionality of the dataset, we performed latent semantic indexing followed by singular value decomposition (Methods). Batch correction was performed using the deep neural network-based scAlign^29^ to correct for technical sources of variance, including individual variation and processing method (Extended Data Fig. 1e-f, Methods). We identified 25 distinct clusters using the Leiden community detection algorithm (Fig. 1c, Extended Data Fig. 1g-h). This analysis robustly separated cortical and subcortical cells (MGE)(Fig. 1d).

Next, to determine which epigenomic signatures correspond to the known cell types of the developing cortex, we calculated ‘gene activity scores’ by summing fragments in the gene body and promoter regions, which represents a proxy for gene expression^20, 30^ (Fig. 1e). Activity of canonical marker genes identified the major cell classes, including radial glia (RG), intermediate progenitor cells (IPCs), excitatory neurons (ENs), and interneurons (INs)(Fig. 1g, Extended Data Fig. 2a-b). To systematically predict cell identity, we correlated gene activity scores from scATACseq cells with cell type marker genes inferred from previously published scRNAseq data^1^ (Methods), and assigned putative cell identity to every cell in the scATACseq dataset according to the highest correlation of scRNAseq-based cluster (Fig. 1f, Extended Data Fig. 2c). Most scATACseq clusters had one-to-one mapping to scRNAseq clusters, with few exceptions. Radial glia formed a single cluster in scATACseq, but map to three scRNAseq clusters of radial glia (‘ventricular’, ‘outer’, and ‘truncated’). Conversely, many clusters of excitatory neurons mapped to a single cluster of PFC or V1 excitatory neurons from scRNAseq (Fig. 1h). These discrepancies suggest that for some cell classes, epigenomic information may provide additional resolution of cell types beyond transcriptional definitions, while in other cases, transcriptomics may reveal more subtypes.

**Fig. 2:**
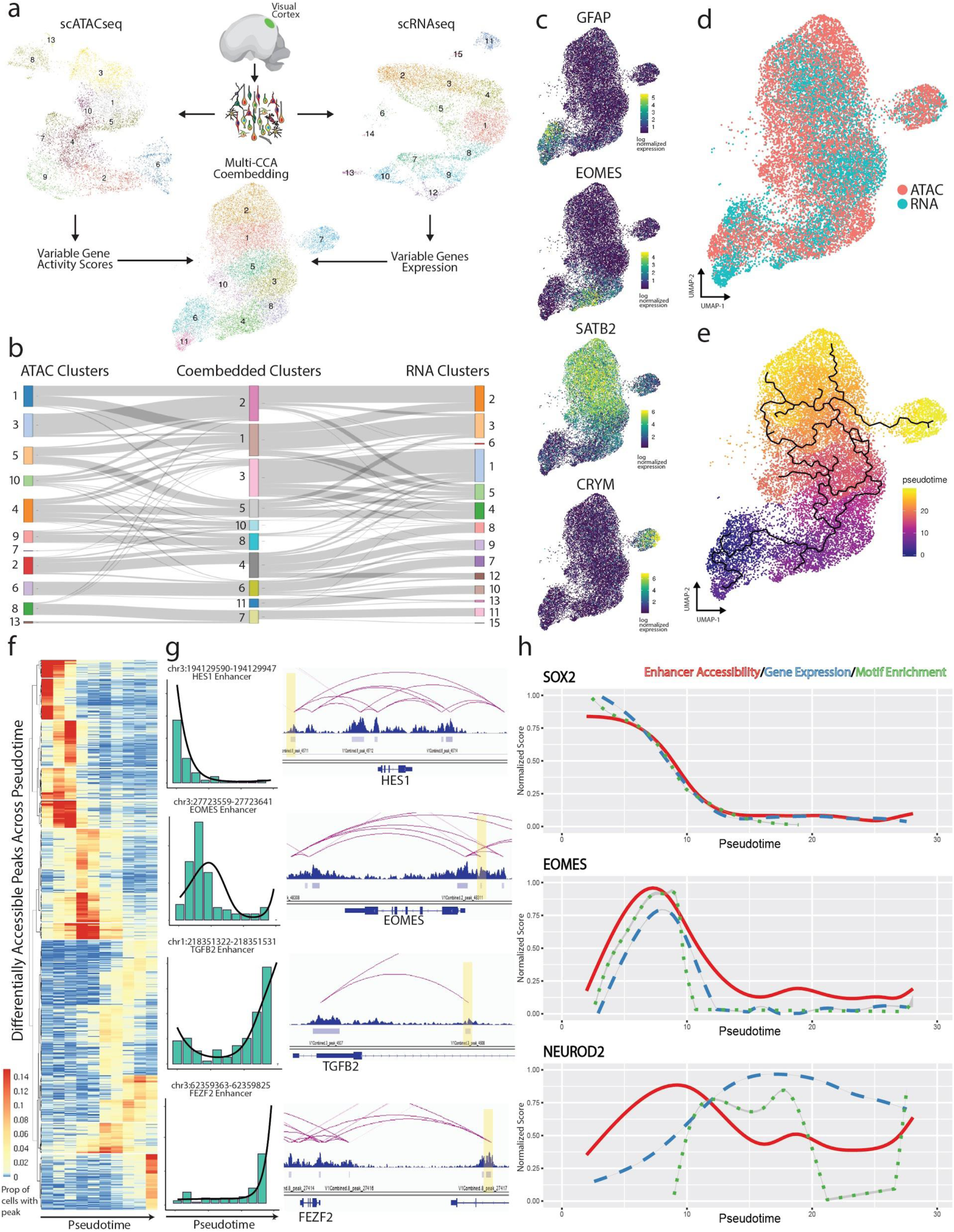
Dynamic changes in chromatin accessibility during human cortical neurogenesis. **a)** Schematic depicting experimental workflow for coembeddding scATACseq and scRNAseq data from the same samples. Top left, UMAP projection of scATAC-seq cells from samples of visual cortex (n = 3 individuals) colored by leiden clusters. Top middle, schematic depicting experimental workflow. Top right, UMAP projection of scRNA-seq cell from samples of visual cortex (n = 2 individuals, same as 2 of 3 from scATACseq) colored by leiden clusters. Bottom middle, UMAP projection of coembedded scATAC-seq & scRNA-seq cells colored by leiden clusters. **b)** Sankey plot depicting the mappings between scATAC-seq clusters, scRNA-seq clusters, and coembedded clusters. **c)** Projection of log normalized gene expression and gene activity scores in coembedded space for GFAP (RGs), EOMES (IPCs), SATB2 (Upper-layer ENs), and CRYM (Deep-layer ENs). **d)** UMAP projection of coembedded cells colored by assay. **e)** UMAP projection of coembedded cells colored by pseudotime with principal graph overlaid. **f)** Heatmap depicting the average proportion of cells with peaks that are differentially accessible across pseudotime. Cells are binned by pseudotime into equally sized bins. **g)** Left, barplots of peak accessibility for 4 individual peaks across 10 pseudotime bins with regression line overlaid. Right, predicted enhancer-gene interaction predictions for each of the four peaks, peaks highlighted in yellow. **h)** Comparison of moving averages of normalized enhancer accessibility (red), gene expression (blue), and motif enrichment (green) across pseudotime for SOX2 (top), EOMES (middle), and NEUROD2 (bottom).

### Single cell chromatin state profiling reveals candidate cell type specific enhancers

To identify putative cell type specific gene regulatory elements, we called peaks on aggregate single cells from each cluster^31^ (Methods, Fig. 1b). Non-overlapping peaks were subsequently merged to a total union set of 398,139 peaks. Cluster-specific differentially accessible peaks were identified for each cluster, resulting in a set of >200,000 DA peaks, with most clusters containing many thousands of cluster specific peaks (Fisher’s Exact, FDR<0.05, Fig. 1h, Extended Data Fig. 3, Extended Data Fig. 4j). Annotation of our peak set in genomic features shows enrichment in intronic and distal intergenic regions and in the flanking regions of transcription start sites, suggesting an enrichment of gene regulatory elements, such as enhancers (Extended Data Fig. 4a-b). We intersected our scATACseq peaks with publicly available ChIP-seq data for H3K27ac (GEO: GSE63648), a marker for active enhancers, generated for comparable tissue samples, and found significant overlap with our peaks (Permutation test, one-sided, p<0.001, Extended Data Fig. 4c). We also intersected our peak set with a set of validated forebrain enhancers^32^ (VISTA Enhancer Browser) and found that 297/317 overlapped with our peakset (Extended Data Fig. 4d). Due to growing evidence that regions of the genome that have undergone accelerated sequence evolution in humans are enriched for neurodevelopmental enhancers^24^, we intersected our peak set with a set of 2,540 non-coding human accelerated regions (ncHARs) finding 880 overlaps (Extended Data Fig. 4e). Interestingly, chromatin accessibility profiles of MGE progenitors and MGE-derived cortical interneurons were enriched for accessibility of ncHARs, and future studies are needed to elucidate if those genomic changes could have contributed to the changes in interneuron repertoire across primates^33, 34^) (Extended Data Fig. 4f-i).

**Fig. 3:**
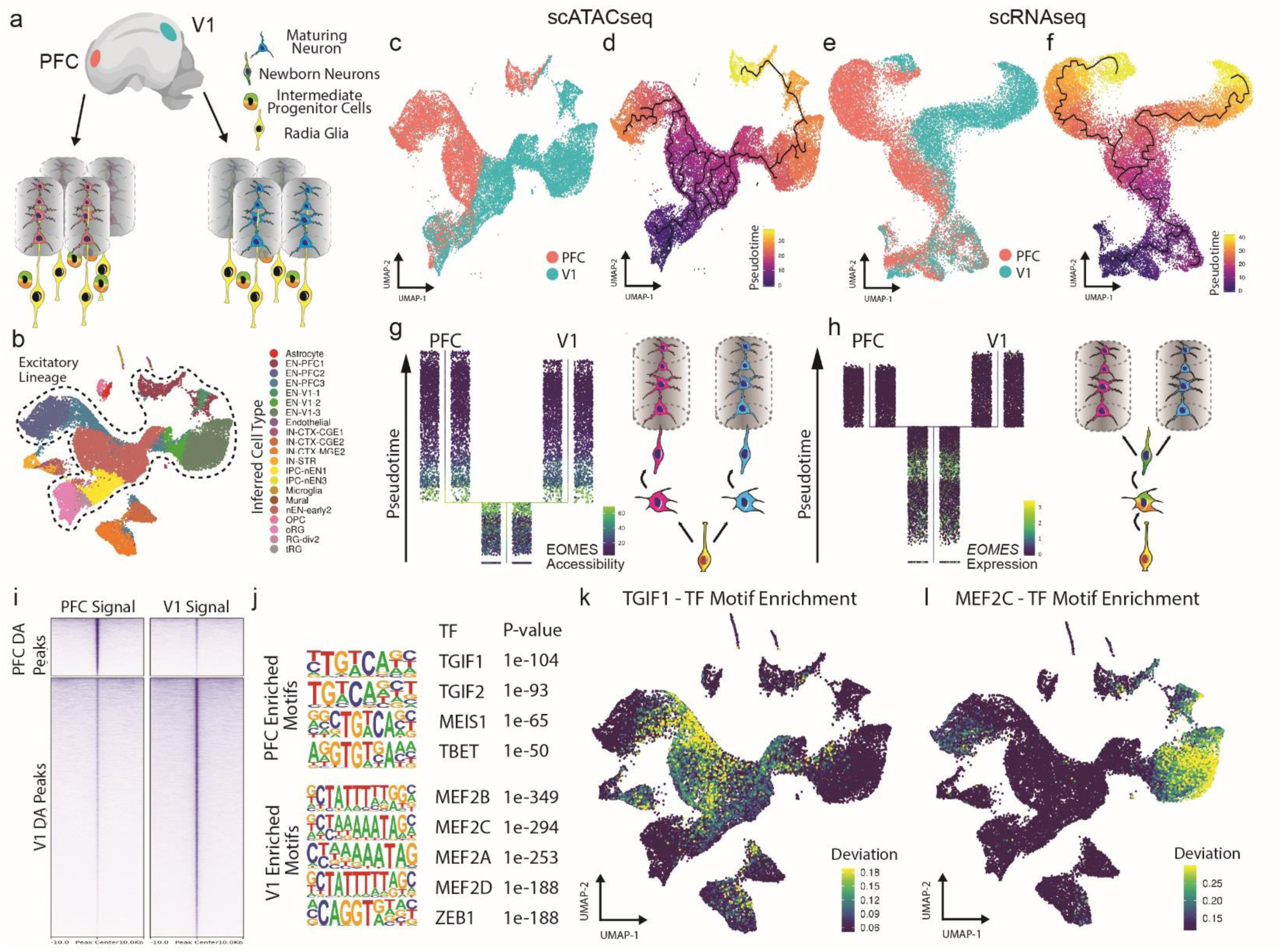
Areal differences in chromatin state of progenitor cells foreshadow the emergence of area-specific excitatory neuron types. **a**) Schematic depicting differentiation trajectories for excitatory neurons from the PFC (left) and V1 (right). **b)** UMAP projection of all PFC & V1 scATACseq cells (n = 3 individuals) colored by cell type predictions. Cells from the excitatory lineage are circled by a dashed line. **c)** UMAP projection of PFC & V1 scATACseq excitatory lineage cells colored by area of origin. **d)** UMAP projection of PFC & V1 scATACseq excitatory lineage cells colored by pseudotime with principal graph overlaid. **e)** UMAP projection of PFC & V1 scRNAseq excitatory lineage cells (n = 2 individuals, same as 2 of 3 from scATACseq) colored by area of origin. **f)** UMAP projection of all PFC & V1 scRNAseq excitatory lineage cells colored by pseudotime with principal graph overlaid. **g)** Left, Projection of PFC & V1 scATACseq excitatory lineage cells ordered from bottom to top by pseudotime value with PFC/V1 divergence branch point displayed (Methods). Cells colored by gene activity score of EOMES, highlighting IPCs. Right, schematic illustrating the excitatory neuron differentiation trajectory based on chromatin accessibility, with which PFC/V1 divergence becomes apparent at the level of IPCs. **h)** Left, Projection of PFC & V1 scRNAseq excitatory lineage cells ordered from bottom to top by pseudotime value with PFC/V1 divergence branch point displayed (Methods). Cells colored by expression of EOMES, highlighting IPCs. Right, schematic illustrating the excitatory neuron differentiation trajectory based on gene expression, with which PFC/V1 divergence is not apparent until the level of maturing neurons. **i)** Pileups of PFC and V1 signal in PFC and V1 specific peak sets. Pileups are centered on peaks showing +/-10Kb flanking regions. **j)** Top, TF motif enrichments in set of 5,863 PFC specific peaks (Fisher’s Exact, two-sided, FDR<0.05) as determined by HOMER. Bottom, TF motif enrichments in set of 26,520 V1 specific peaks (Fisher’s Exact, two-sided, FDR<0.05) as determined by HOMER. **k)** UMAP projection of deviation scores of motif enrichment for TGIF1 as determined by ChromVAR. **l)** UMAP projection of deviation scores of motif enrichment for MEF2C as determined by ChromVAR.

**Fig. 4:**
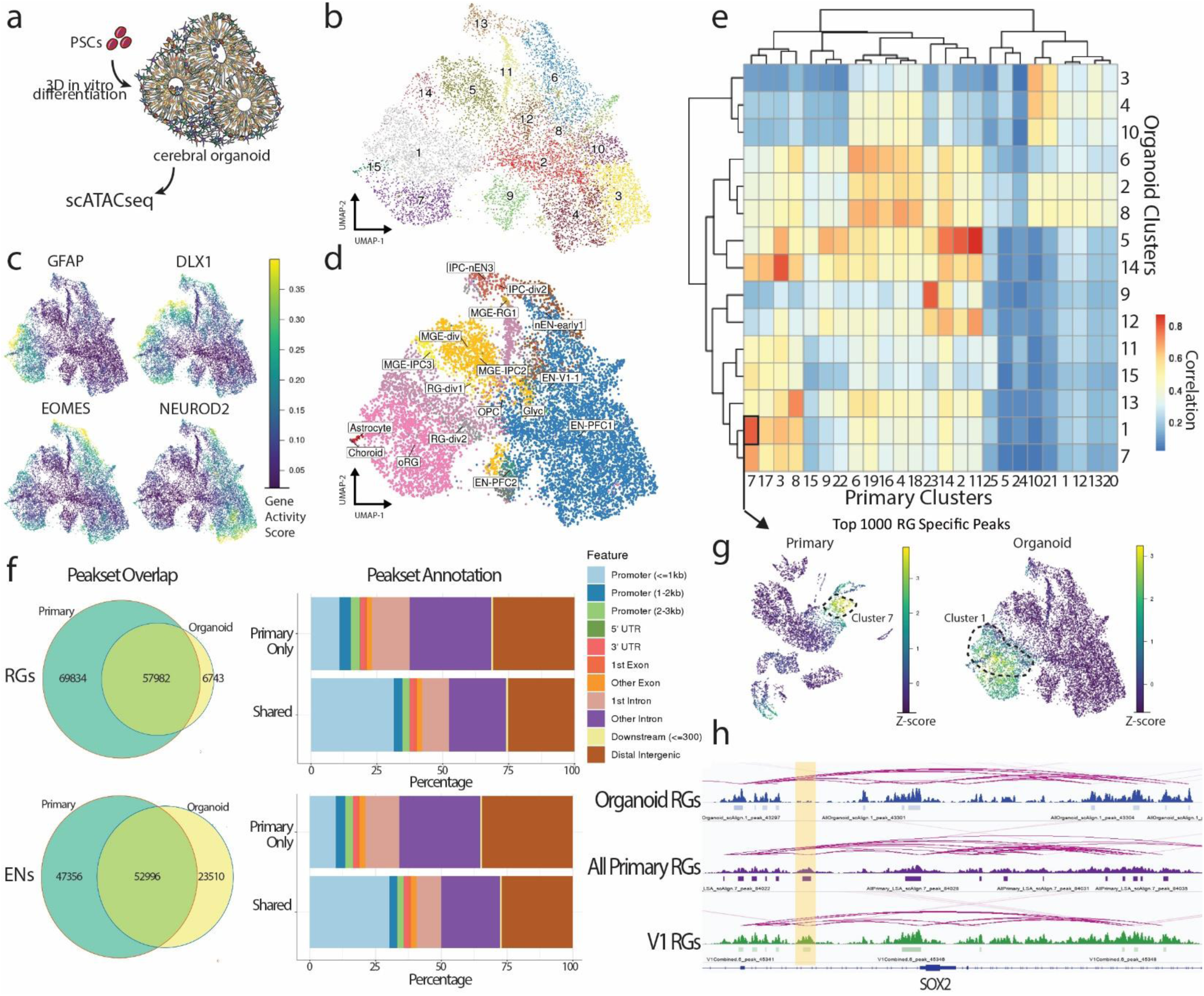
Cell type-specific differences in chromatin accessibility between cerebral organoids and the developing human brain. **a)** Schematic depicting experimental workflow. PSCs – pluripotent stem cells. **b)** UMAP projection of all organoid scATACseq cells (n = 3 organoids from different lines, 11,171 cells) colored by leiden clusters. **c)** UMAP projections of gene activity scores for GFAP marking radial glia, EOMES marking intermediate progenitors, DLX1 marking interneurons, and NEUROD2 marking excitatory neurons. **d)** UMAP projection of all organoid scATACseq cells colored by cell type predictions (Methods). **e)** Heatmap of pearson correlations between primary and organoid scATACseq clusters based on a common peak set. **f)** Top left, overlap of primary and organoid radial glia peak sets. Top right, annotation of primary only RG peaks and shared RG peaks in genomic features. Bottom left, overlap of primary and organoid excitatory neuron peaksets. Bottom right, annotation of primary only EN peaks and shared EN peaks in genomic features. **g)** Left, UMAP projection of enrichment Z-scores of the top 1000 radial glia specific peaks (Fisher’s Exact, two-sided) in all primary scATACseq cells. Right, UMAP projection of Z-scores of enrichment of the same 1000 radial glia specific peaks in all organoid scATACseq cells. **h)** Genome browser tracks of the SOX2 locus showing enhancer-gene predictions for organoid radial glia (top, blue), all primary radial glia (middle, purple), and V1 radial glia (bottom, green). Highlighted in yellow is a peak that is predicted to interact with SOX2 present in both primary radial glia populations and not present in the organoid radial glia.

To identify putative regulatory ‘grammar’ of cell types, we calculated enrichment of known transcription factor binding motifs in cluster specific peak sets using HOMER^35^ (Methods, Fig. 1h). Transcription factor motif enrichments align with cell type annotations from marker gene body enrichments with NEUROD1 motif enrichment in EN clusters, DLX and ASCL1 motif enrichments in IN clusters, PAX6 motif enrichment in RG clusters, EOMES motif enrichment in IPC clusters, SOX9 motif enrichment in OPC clusters, IRF8 motif enrichment in microglia clusters, and NKX2.1 motif enrichment in MGE progenitors. To examine transcription factor motif enrichments at the single cell level, we used ChromVAR, which estimates bias-corrected deviations of transcription factor binding motif enrichments in scATAC-seq libraries, and found good agreement with top motif enrichments for each cluster as determined by HOMER (Extended Data Fig. 4k). Finally, we predicted likely enhancers using the recently developed activity-by-contact (ABC) model^36^, which integrates H3K27ac ChIP-seq, Hi-C, and gene expression data with chromatin accessibility to predict enhancers and link them to their target genes (Methods). Using this method, we were able to identify sets of high-confidence putative enhancers for each cell type and their likely target genes (Fig. 1i).

### Vulnerability of cell type specific regulatory landscape to neurodevelopmental disorders

Mutations in non-coding genomic regions, as well as *de novo* loss of function mutations in chromatin regulators have been implicated in a wide range of neurodevelopmental and psychiatric disorders, including schizophrenia^37^ and autism spectrum disorder^38–41^. However, due to the lack of cellular-resolution datasets of chromatin state across developmental stages and differentiation states, these mutations cannot be tied to selective vulnerabilities across diverse cell types of the developing human brain. To address this unmet need, we intersected cell type specific ATAC-seq peaks with disease-linked common and rare non-coding variants (Methods). We first intersected our cell type-specific peak sets with *de novo* non-coding mutations (DNMs) identified from ASD and neurodevelopmental delay (NDD) cases and found significant enrichment of DNMs in 19 of 27 cell-type specific peak sets, compared to a merged background peak set (Extended Data Figure 5). However, no cell type-specific peak sets were significantly enriched for DNMs in probands compared to sibling controls. We also intersected cell type specific peak sets with genomic regions enriched for copy number variants in cases with developmental delay^42^, identifying cell types with significant enrichment and depletion (Figure 1j). Because such regions are large and do not provide specificity with respect to individual genes, we next tested for enrichment of cell type specific peaks in the flanking regions of genes associated with ASD and NDD and identified cell types with peak sets significantly enriched and depleted in these regions (Figure 1k), but were underpowered to identify any differences in the DNM burden in peaks within the promoter and gene body between probands and siblings across peak sets. Finally, we sought to assess the enrichment of common variants associated with neuropsychiatric disease risk in our cell type specific peak sets. To do this we performed a partitioned heritability LD score regression analysis using summary statistics from large-scale genome-wide association studies of schizophrenia, ASD, major depressive disorder, and bipolar disorder (Methods). For all four disorders, we detected significant enrichments (FDR < 0.05) of risk-associated variants in peaks from at least one cell type. Interestingly, we found that, consistently across disorders, disease risk was most strongly enriched in excitatory neuron populations (Figure 1l).

### Dynamic changes in chromatin accessibility during neuronal differentiation

Developing tissues pose unique challenges to single cell analysis methods because, unlike the adult tissue, many cells represent developmentally transient states along the continuum of lineage progression. Chromatin state profiling provides a unique opportunity to characterize the Waddingtonian landscape of cell fate decisions underlying the emergence of cell types during development. Identification of putative regulatory mechanisms of cell fate specification could in turn be harnessed to promote directed differentiation of molecularly-defined cell types from pluripotent stem cells for applications in cell replacement therapy and disease modelling. To combine transcriptomic and epigenomic information, we coembedded scRNAseq and scATACseq datasets generated in parallel from samples of visual cortex (Fig. 2a). By projecting cluster annotations across the three comparisons (scRNAseq only, scATACseq only, and multimodal mapping), we were able to further support our predictions of cellular classification from chromatin state data (Fig. 2b). Projections of gene expression and gene activity scores in the co-embedded space show that distinct clustering of unique cell types is preserved and that scATACseq and scRNAseq cells of the same type cluster together (Fig. 2c-d). Because the ability to identify cell types across samples, species, and experimental models using single cell genomics approaches depends on robust detection of gene co-expression relationships^1, 43^, we sought to compare gene-module assignments calculated based on mRNA expression values detected using scRNAseq or those inferred from scATACseq. We combined genes into modules using weighted gene co-expression network analysis^44^ and compared eigengene projections as well as gene-module assignments between scRNAseq and scATACseq datasets (Extended Data Fig. 6a-b). This analysis revealed a remarkable conservation of gene co-expression relationships, with the exception of genes related to signaling pathway activation that formed co-expression module in scRNAseq but not scATACseq datasets, and genes related to synapse assembly forming co-expression module in scATACseq, but not scATACseq dataset (Extended Data Fig. 6d-g). Together, our joint analysis of scRNAseq and scATACseq datasets further underscored the conserved representation of the major cell types and gene co-expression relationships across the two modalities.

To identify trajectories of chromatin accessibility underlying excitatory neuron differentiation and maturation, we performed pseudotemporal ordering of cells in the co-embedded space^45^ (Fig. 2e, Methods). Consistent with the known patterns of neurogenesis, pseudotime reconstruction ordered sequentially radial glia, intermediate progenitors, and excitatory neurons. We identified hundreds of loci with sharp, transient accessibility across pseudotime, and predicted enhancers that interact with genes linked to cell type identity (Fig. 2f,g). By calculating transcription factor binding site enrichment across peaks that show dynamic changes in accessibility along pseudotime, we reconstructed the known hierarchy of transcription factors involved in cortical neurogenesis, including sequential enrichment for SOX2, ASCL1, and NEUROD2 binding sites among transiently accessible loci (Extended Data Fig. 7). These results challenge the prevailing model of differentiation as a transition between two phases involving progressive loss of accessibility of sites open in progenitor cells and gradual opening of sites relevant to postmitotic cells^46^, and underscore highly dynamic transient states of chromatin accessibility during human cortical neurogenesis.

Furthermore, we leveraged the scRNAseq and scATACseq co-embedding to compare changes in gene expression, enhancer accessibility, and transcription factor motif enrichment along the differentiation trajectory. Considering a few key regulators of neurogenesis, SOX2, EOMES, and NEUROD2, we observed a trend for accessibility of predicted enhancers to precede changes in gene expression (Fig. 2h). These findings are consistent with recent reports^21, 47^ and support the model whereby changes in chromatin state along a developmental lineage foreshadow changes in gene expression and cell fate decisions. Intersection of cell type and developmentally dynamic loci and putative regulatory elements with whole genome sequencing data from neurodevelopmental or neuropsychiatric disorders may reveal developmentally transient states that are vulnerable to non-coding mutations.

### Cortical progenitors develop area-specific chromatin states

Single cell transcriptomics recently revealed that area-specific cortical excitatory neurons emerge during early neurogenesis, while only limited molecular differences can be found between progenitor cells^1, 2^. Given that changes in the accessibility of regulatory elements often precede changes in gene expression (Fig. 2h), we sought to examine whether epigenomic signatures could foreshadow the emergence of area-specific excitatory neurons. Specifically, we compared scRNAseq and scATACseq profiles of excitatory lineage cells sampled from the extremes of the rostral-caudal axis, PFC and V1 (Fig. 3a-b, Extended Data Fig. 8a-h). For each modality, we ordered the cells in pseudotime to approximate the differentiation trajectory, and identified the ‘branch’ point along this trajectory at which transcriptomic or chromatin state differences between PFC and V1 lineages become apparent. In contrast to transcriptomic data, which only distinguishes maturing excitatory neuron clusters from distinct cortical areas^1^ (Fig. 3h), chromatin state signatures reveal a striking divergence between PFC and V1 earlier in differentiation, and define area-specific IPC populations (Fig. 3g).

To identify putative regulatory networks that could underlie the divergence of PFC and V1 lineages, we performed transcription factor binding site enrichment analysis^35^ on peaks that were differentially accessible between PFC and V1 (Fisher’s Exact, two-sided, FDR<0.05, Fig. 3i-l, Extended Data Figure 8i-j, Supplementary Tables 1&2). This analysis recovered previously known regulators of cortical arealization and those consistent with transcriptomic studies^1, 48^. For example, our analysis predicts motif enrichment for known transcription factors enriched in the PFC, including POU3F2, MEIS1, TBR1, NEUROD1, NEUROG2, and TBX21 (Supplementary Table 1). Interestingly, many of the candidates identified in this analysis relate to retinoic acid signaling pathway. In early development, retinoic acid signaling plays a well-established role in development of caudal fates, including hindbrain and spinal cord^49^. However, at later stages of development, retinoic acid signaling has been shown to interact with pathways involved in cortical arealization including the NR2F1 transcription factor and Wnt signaling that promote occipital (visual cortex) identities^50, 51^, and is negatively regulated by TGIF1^52^, the top enriched motif among PFC cells (Fig. 3j-k). Together, our analyses suggest that epigenomic differences distinguish cortical progenitor cells between cortical areas and foreshadow the emergence of area-specific excitatory neuron subtypes. In addition, our study suggests a previously unappreciated role for the retinoic signaling pathway in cortical arealization.

### An epigenomic ‘report card’ for in vitro models of cortical neurogenesis

Due to the scarcity of primary human tissue, studies of human neural development critically require suitable *in vitro* models. Cerebral organoids are a three-dimensional culture model of the developing brain that can be derived from somatic cells. Previous studies emphasized the similarities between cerebral organoid cells and their *in vivo* counterparts using single cell transcriptomics^43, 53, 54^ and bulk epigenomics^15, 17^. We sought to extend these comparisons by performing chromatin state profiling of cerebral organoids at single cell resolution and generated scATACseq data for cortical organoids derived via directed differentiation from three genetically normal individuals^43^ (Fig. 4a, Extended Data Fig. 9a, Methods). We identified the major classes of cell types expected to emerge in this model, including radial glia, IPCs, interneurons, and excitatory neurons, although individual clusters were less discrete than their in vivo counterparts, and contained fewer distinguishing chromatin state features (Fig. 4b-d, Extended Data Fig. 9d,f).

To compare organoid clusters with their primary counterparts, we quantified the chromatin accessibility signal from organoid cells in peaks defined from primary cells, allowing us to identify clusters representing homologous cell types (Methods, Fig. 4e). We found that cell type specific peaks identified in primary cells maintained cell type specificity in organoid cells, but many peaks corresponding to cell types not present in organoids, such as microglia and endothelial cells, were missing (Fig. 4g, Extended Data Fig. 9e). Next, to assess the fidelity of organoids as a model for the epigenomic state of primary cortical cells, we called peaks for each organoid cluster and compared with primary peaks for homologous cell types. We found that, while organoids mostly contain peaks found in their corresponding primary counterparts, they are missing ∼50% of peaks identified in primary clusters (Fig. 4f, Extended Data Fig. 9b-c). Interestingly, shared peaks show higher enrichment in promoter regions, while peaks found only in primary are more enriched in distal intergenic and intronic regions, suggesting that organoids may be missing many distal regulatory elements identified in primary cells. In addition, we compared enhancers predicted by activity-by-contact model, which revealed that organoids lack many candidate cell type specific enhancers found in primary, even after correcting for cellular coverage (Fig. 4h).

In summary, scATACseq data generated for primary cells represents a blueprint for normal epigenomic states of cell types in the developing human brain that serves as a reverence for evaluating the fidelity and robustness of *in vitro* derived models. Our analysis reveals features of chromatin state found in normal developing brain are recapitulated in cerebral organoids, epigenomic features of organoid cell types are less discrete and lack thousands of distal regulatory elements found *in vivo*.

## Discussion

By performing massively parallel single cell profiling of chromatin state, we were able to extend previous studies of cell-type specific epigenomic regulation of brain development. Specifically, scATAC-seq analyses reveals transiently accessible loci that track with neuronal differentiation. These states may reveal mechanisms governing the establishment of cell fate during neurogenesis, in particular those underlying the emergence of area-specific excitatory neurons. Our findings provide new perspectives on the mechanisms underlying the rewiring of regulatory pathways during neurogenesis^46^. Intersection of chromatin state landscape with disease variants implicates post-mitotic, developing cortical excitatory neurons in the etiopathogenesis of neuropsychiatric disorders^14, 55, 56^, and future studies are needed to probe how disease-associated variants in these regulatory regions modify cell fate decisions in the developing cortex. Furthermore, our findings suggest a potential limitation for cortical organoids to serve as experimental models to test these hypotheses, as many non-coding regulatory elements, in particular distal enhancers, may not be recapitulated in this model. In order to determine the fidelity of organoids as a model for the epigenomic landscape of the developing brain, functional characterization of conserved and non-conserved chromatin features *in vivo* and *in vitro* will be required. In addition to serving as a reference for evaluating *in vitro* models of human brain development, our data will enable scalable prediction of candidate cell type specific regulatory elements that could facilitate the development of enhancer-based tools for accessing molecularly-defined cell types^57–59^.

## Author Contributions

R.S.Z. and T.J.N. designed the experiments. R.S.Z. performed all experiments. R.S.Z. and C.N.K. performed the majority of data analysis. P.F.P. and K.S.P. contributed to the analysis. A.W., T.N.T., A.K., A.M.C., S.A.A., and E.E.E. performed the disease intersection analyses. R.S.Z., C.N.K., N.A, and T.J.N. wrote the paper with input from all authors. M.H. built the cell browser interface. T.J.N. and N.A. supervised the project.

## Acknowledgements

We thank Aparna Bhaduri for help with generation of scRNAseq data and its analysis, and helpful discussions throughout the project, as well as Madeline Andrews for sharing organoid cultures. We thank John Rubenstein for reading of the manuscript. This study was supported by NIH awards: psychENCODE award U01MH116438, Brain Initiative award U01MH114825, and the Psychiatric Cell Map Initiative Convergence Neuroscience award U01MH115747, Autism Speaks Predoctoral Fellowship (11874 to R.S.Z.), NARSAD Young Investigator Grant (to T.J.N), gifts from Schmidt Futures and the William K. Bowes Jr. Foundation, and the National Institute of Mental Health (1K99MH117165 to T.N.T.).

## Data Availability

scRNA-seq of the PFC/V1 will be available on the NeMo archive (accession numbers pending).

## METHODS

### Tissue Source

De-identified tissue samples were collected with previous patient consent in strict observance of the legal and institutional ethical regulations. Protocols were approved by the Human Gamete, Embryo, and Stem Cell Research Committee (institutional review board) at the University of California, San Francisco.

### Nuclei isolation from fresh primary tissue

Cortical areas were microdissected from 3 specimens of mid-gestation human cortex, in addition to 3 specimens of non-area-specific mid-gestation human cortex. Tissue was dissociated in Papain containing Deoxyribonuclease I (DNase) for 30 minutes at 37C and samples were triturated to form a single cell suspension. 10^6^ Cells were pelleted and lysed for 3 minutes in 100uL chilled Lysis Buffer (10mM Tris-HCl pH7.4, 10mM NaCl, 3mM MgCl2, 0.1% Tween-20, 0.1% Igepal CA-630, 0.01% Digitonin, 1% BSA). Lysed cells were then washed with 1mL chilled Wash Buffer (10mM Tris-HCl pH7.4, 10mM NaCl, 3mM MgCl2, 0.1% Tween-20, 1% BSA) and nuclei were pelleted at 500rcf for 5 minutes at 4C.

### Nuclei isolation from frozen primary tissue

Tissue sections were snap frozen and stored at -80C. Nuclei were isolated from frozen tissues using the protocol published in Corces MR et al., 2017^1^. Briefly, frozen tissue samples were thawed in 2mL chilled Homogenization Buffer (10mM Tris pH7.8, 5mM CaCl2, 3mM Mg(Ac)2, 320 mM Sucrose, 0.1mM EDTA, 0.1% NP40, 167uM β-mercaptoethanol, 16.7uM PMSF) and lysed in a pre-chilled dounce. Cell lysates were then centrifuged in an Iodixanol gradient for 20 minutes at 3000rcf at 4C in a swinging bucket centrifuge with the brake turned off. The nuclei band was then carefully pipetted and nuclei were diluted in Wash Buffer.

### Cortical organoid differentiation protocol

Cortical organoids were cultured using a forebrain directed differentiation protocol^2, 3^. Briefly, 3 genetically normal PSC lines, H28126, 1323-4, and H1 (WA01), were expanded and dissociated to single cells using accutase. After dissociation, cells were reconstituted in neural induction media at a density of 10,000 cells per well in 96 well v-bottom low adhesion plates. GMEM-based neural induction media includes 20% Knockout Serum Replacer (KSR), 1X non-essential amino acids, 0.11mg/mL Sodium Pyruvate, 1X Penicillin-Streptomycin, 0.1mM Beta Mercaptoethanol, 5uM SB431542 and 3uM IWR1-endo. Media was supplemented with 20uM Rock inhibitor Y-27632 for the first 6 days. After 18 days organoids were transferred from 96 to six well low adhesion plates and moved to an orbital shaker rotating at 90rpm and changed to DMEM/F12-based media containing 1X Glutamax, 1X N2, 1X CD Lipid Concentrate and 1X Penicillin-Streptomycin. At 35 days, organoids were moved into DMEM/F12-based media containing 10% FBS, 5ug/mL Heparin, 1X N2, 1X CD Lipid Concentrate and 0.5% Matrigel. Throughout culture duration organoids were fed every other day.

### Nuclei isolation from cerebral organoids

Cerebral organoids were dissociated in Papain containing Deoxyribonuclease I (DNase) for 30 minutes at 37C and samples were triturated to form a single cell suspension. 10^6^ Cells were pelleted and lysed for 3 minutes in 100uL chilled Lysis Buffer (10mM Tris-HCl pH7.4, 10mM NaCl, 3mM MgCl2, 0.1% Tween-20, 0.1% Igepal CA-630, 0.01% Digitonin, 1% BSA). Lysed cells were then washed with 1mL chilled Wash Buffer (10mM Tris-HCl pH7.4, 10mM NaCl, 3mM MgCl2, 0.1% Tween-20, 1% BSA) and nuclei were pelleted at 500rcf for 5 minutes at 4C.

### Single Cell RNA-seq Library Preparation and Sequencing

Single cell RNA-seq libraries were generated using the 10x Genomics Chromium 3’ Gene Expression Kit. Briefly, single cells were loaded onto chromium chips with a capture target of 10,000 cells per sample. Libraries were prepped following the provided protocol and sequenced on an Illumina NovaSeq with a targeted sequencing depth of 50,000 reads per cell. BCL files from sequencing were then used as inputs to the 10X Genomics Cell Ranger pipeline.

### Single Cell RNA-seq Analysis

For preprocessing of scRNA-seq data, a minimum of 500 genes and 5% mitochondrial cutoff was used and Scrublet^4^ for doublet removal. The SCTransform^5^ workflow in Seurat^6^ were run separately on each batch.

Canonical component analysis (CCA) on the Pearson residuals from SCTransform was used as input into scAlign for batch correction.

### Bulk ATAC-seq Library Preparation and Sequencing

Bulk ATAC-seq libraries were generated using the protocol outlined in Corces MR et al., 2017 (^1^). Briefly, 50,000 nuclei were permeablized and tagmented. Tagmented chromatin libraries were generated and sequenced on an Illumina NovaSeq with a target sequencing depth of 50 million reads per library. Sequencing data was used as an input to the ENCODE ATAC-seq analysis pipeline (https://github.com/ENCODE-DCC/atac-seq-pipeline).

### Single Cell ATAC-seq Library Preparation and Sequencing

Nuclei were prepared as outlined in the 10X Genomics Chromium single cell ATAC-seq solution protocol. Nuclei were loaded with a capture target of 10,000 nuclei per sample. scATAC-seq libraries were prepared for sequencing following the 10X Genomics single cell ATAC-seq solution protocol. scATAC-seq libraries were sequenced using PE150 sequencing on an Illumina NovaSeq with a target depth of 25,000 reads per nucleus (see Extended Data Table 1).

### Single Cell ATAC-seq Analysis Pipeline

#### Cell Ranger

BCL files generated from sequencing were used as inputs to the 10X Genomics Cell Ranger ATAC pipeline. Briefly, FASTQ files were generated and aligned to GRCh38 using BWA. Fragment files were generated containing all unique properly paired and aligned fragments with MAPQ > 30. Each unique fragment is associated with a single cell barcode.

#### SnapATAC

Fragment files generated from the Cell Ranger ATAC pipeline were loaded into the SnapATAC^7^ pipeline (https://github.com/r3fang/SnapATAC) and Snap files were generated. A cell-by-bin matrix was then generated for each sample by segmenting the genome into 5-Kb windows and scoring each cell for reads in each window. Cells were filtered based on log10(UMI) between 3-5 and fraction of reads in promoters between 10-60% to obtain cells with high quality libraries. Bins were then filtered, removing bins overlapping ENCODE blacklist regions (http://mitra.stanford.edu/kundaje/akundaje/release/blacklists/). This matrix was then binarized and coverage of each bin was calculated and normalized by log10(count + 1). Z-scores were calculated from normalized bin coverages and bins with a z-score beyond ± 2 were filtered from further analysis. A cell-by-cell similarity matrix was generated by calculating the Latent Semantic Index (LSI) of the binarized bin matrix. Principal component analysis (PCA) was then performed on LSI values. The top 50 principal components were used for batch correction through scAlign.

#### scAlign Batch Correction

Multiple batches were integrated using the scAlign package^8^ (https://github.com/quon-titative-biology/scAlign). The ATAC batches were first merged together to calculate the Latent Semantic Index (LSI) with the TF matrix log-scaled for input into PCA. The 50 principal components of LSI were used as inputs to the encoder. The latent dimension was set at 32 and ran with all-pairs alignment of all batches. The input dimension to the encoder was set to 50 to match the input 50 principal components, and trained to 15,000 iterations using the small architecture setting with batch normalization (BN). The 32 dimensions were used for downstream analysis for finding neighbors. The scRNAseq were processed using Seurat and computed the top 15 components from CCA for input into scAlign, and the latent dimension was set to 20 using the small architecture with BN and 15,000 iterations. All alignments were unsupervised.

#### Clustering and Visualization

In order to visualize the high dimensionality dataset in 2D space, the latent dimensions for the ATAC and RNA data from scAlign were used to construct UMAP (https://arxiv.org/abs/1802.03426) graphs from Seurat. A K-nearest neighbor graph was constructed using the latent dimensions from scAlign. The leiden algorithm was then used to identify ‘communities’, or clusters, in the sample, representing groups of cells likely to be of the same cell type.

#### Calculating Gene Activity Scores

To create a proxy for gene expression, ATACseq fragments in the gene-body plus promoter (2Kb upstream from transcription start sites) of all protein-coding genes were summed for each cell to generate ‘Gene Activity Scores’. A matrix was constructed for all gene activity scores by all cells. Due to the sparsity of scATAC-seq data, the MAGIC imputation method^9^ was used to impute gene activity scores based on the K-nearest neighbor graph.

#### Cell Type Predictions

In order to link scATACseq clusters to known cell types of the developing cortex, gene activity scores were used to correlate scATACseq cells with cell types from scRNAseq data generated from similar samples^10^. Briefly, cluster ‘marker genes’ were identified for each leiden cluster by performing a Fisher’s Exact test for enrichment of gene activity scores. The top 300 enriched genes for each cluster were identified based on p-value. Similarly, cell type marker genes were identified for annotated cell types from Nowakowski et al. 2017^10^, and the top 300 genes for each cluster were identified based on p-value. The intersection of genes from scATACseq marker genes and scRNAseq marker genes was identified, resulting in a set of 1084 genes. Average expression values for each scRNAseq cell type for each of the 1084 genes was determined. For each scATACseq cell, the gene activity scores of the 1084 genes was correlated with cell type averages from the scRNAseq data and a cell type prediction was made based on the cell type most highly correlated with each scATACseq cell.

#### Peak Calling

Fragments from cells were grouped together by cluster and peaks were called on all cluster fragments using MACS2 (https://github.com/taoliu/MACS) with the parameters ‘--nomodel --shift -37 --ext 73 --qval 1e-2 -B -- SPMR --call-summits’. Peaks from each cluster were then combined to form a master peak set and a cell-by-peak matrix was constructed. This matrix was binarized for all downstream applications.

#### Determination of Differentially Accessible Peaks

Differentially accessible peaks for each cluster were determined by performing a two-sided Fisher’s exact test, and selecting peaks that had log fold change >0, and FDR-corrected p-value < 0.05.

#### Visualizing Cluster Signal in Peaks

The deeptools suite^11^ (https://deeptools.readthedocs.io/en/develop/) was used to visualize pileups of cluster-specific ATACseq signal (output from MACS2) in DA peak sets.

#### Transcription Factor Motif Enrichment Analysis

The findMotifsGenome.pl tool from the HOMER suite^12^ (http://homer.ucsd.edu/homer/) was used to identify TF motif enrichments in peak sets. The ChromVAR R package^13^ was used to identify TF motif enrichments at the single cell level in scATACseq data. Briefly, the peak-by-cell matrix from the snap object was used as an input, filtering for peaks open in at least 10 cells. Biased-corrected TF motif deviations were calculated for the set of 1,764 human TF motifs for each cell.

#### Enhancer-Gene Predicted Interactions

The Activity-by-Contact (ABC) model^14^ (https://github.com/broadinstitute/ABC-Enhancer-Gene-Prediction) was used for prediction of enhancer-gene interactions from scATACseq data. Cluster-specific ATAC-seq signal and peak outputs from MACS2 were used as inputs. Gene expression values from scRNAseq of GW20 visual cortex were averaged and used an input. As recommended and provided by the creators of ABC, an averaged Hi-C profile of 10 cell types was used as an input.

#### VISTA Enhancer Intersections

VISTA Enhancers were taken from the VISTA Enhancer Browser (https://enhancer.lbl.gov/) and filtered for human sequences found to be active in the forebrain. Enhancers were lifted over to Hg38 using the UCSC LiftOver tool (https://genome.ucsc.edu/cgi-bin/hgLiftOver) and overlapping regions were merged, resulting in 317 unique regions. These regions were intersected with the peak set from all primary scATACseq cells and 297 peaks overlapping VISTA forebrain enhancer regions were identified.

#### Non-coding Human Accelerated Region Intersections

Non-coding Human Accelerated Regions (ncHARs) were taken from Capra et al. 2013^15^. These regions were then lifted over to Hg38 using the UCSC LiftOver tool and overlapping regions were merged, resulting in 2,540 unique regions. These regions were intersected with the peak set from all primary scATACseq cells and 880 peaks overlapping ncHAR regions were identified.

#### H3K27ac ChIP-seq Data Intersection

Publicly available H3K27ac ChIP-seq peak sets generated from 12pcw (GW14) human cortical samples^16^ were obtained from GEO (GEO: GSE63648). All peak sets were lifted over to Hg38 using the UCSC LiftOver tool and overlapping regions were merged. Using the RegioneR R package^17^ (https://www.bioconductor.org/packages/release/bioc/html/regioneR.html), a one-sided permutation test was performed to determine the significance of overlap between H3K27ac peaks and scATACseq peaks, using 1000 random shuffling iterations to build a null distribution.

#### Genomic Feature Annotations

The ChIPSeeker R package^18^ (https://bioconductor.org/packages/release/bioc/html/ChIPseeker.html) was used to annotate all peak sets in genomic features.

#### Pseudotime Analysis

The Monocle 3 R package^19^ (https://cole-trapnell-lab.github.io/monocle3/) was used for pseudotime calculation of the coembedded RNA and ATAC dataset. The radial glia cells were set as the root cells. The minimum branch length was 9 in the graph building. Monocle 3 was also used for the pseudotime calculation of the scRNAseq PFC/V1 dataset. The Cicero package^20^ (https://cole-trapnell-lab.github.io/cicero-release/) was used for the pseudotime calculation of the scATACseq PFC/V1 dataset.

#### Comparison of Accessibility, Gene Expression, and TF Motif Enrichment Across Pseudotime

Since pseudotime was calculated on the co-embedded space of ATAC and RNA cells, we can directly compare temporal changes in gene expression and chromatin accessibility. For each of the transcription factors, we identified gene-linked enhancers candidates using the ABC model and calculated a 1,000 cell moving average of their accessibility across pseudotime from the ATAC cells. Using Z-scores from ChromVAR, we calculated a 1,000 cell moving average of the motif enrichment across pseudotime from the ATAC cells. For gene expression, we calculated a 1,000 cell moving average across pseudotime from the RNA cells. LOESS regression lines were fit to the moving average data.

#### Branchpoint Analysis

URD^21^ (https://github.com/farrellja/URD/) was used to compare the branchpoint of ATAC and RNA independently. Deep-layer neurons weren’t considered during this analysis due to obfuscating identities, and the batch corrected values were used as input to the diffusion map calculations to combat batch effects. Diffusion parameters were set to 150 nearest neighbors, and sigma was auto calculated from the data. The tree was constructed using 200 cells per pseudotime bin, 6 bins per pseudotime window, and branch point p-value threshold of 0.001.

#### scRNAseq/scATACseq Coembedding

To anchor mRNA expression and chromatin state profiles in the same map of cell diversity, we applied scAlign on datasets where we profiled scRNAseq and scATACseq in parallel in the sample sample. This was achieved by linking gene expression data to gene activity scores derived from chromatin accessibility data. The gene activity scores were logRPM values derived from gene activity scores generated by the SnapATAC pipeline. Then the gene expression and gene activity scores were processed using Seurat, and then split into batches for input into scAlign. The encoder space was computed using multi CCA of the 10 dimensions with latent dimensions at 20 using the ‘small’ architecture.

### Disease Intersection

#### De novo mutation (DNM) enrichment

Peak sets from 27 cell types were intersected DNMs from 2,708 probands and 1,876 siblings using bedtools v2.24.0. Peak sets were determined based on the presence of the peak in at least 5% of the cells for that cell type. DNMs were identified by an in-house pipeline. Briefly, variants from whole-genome sequencing data were called using four independent callers: GATK v3.8, FreeBayes, Strelka, and Platypus. Variant calls from each caller were intersected, and filtered for read depth (> 9), allele balance (> 0.25), absence of reads supporting the mutation in parents, and identified by at least three of the four callers.

Peak sets were tested for an enrichment of DNMs in probands as compared to a background peak set which contained all peaks. We used a Fisher’s exact test to compare the number of peaks with one or more DNMs between the cell type-specific peak set and the background peak set. We also performed a Wilcox rank sum test comparing the number of DNMs per peak in the cell type-specific set to the background peak set. We applied a Bonferroni multiple test correction for 27 tests (# of cell types) to all p-values.

#### ASD/NDD gene set enrichment

We created gene plus promoter regions using bedtools v2.24.0, where we defined the promoter as the 1Mb region upstream of the gene transcription start sites. Gene regions were defined using Gencode V27. Peak sets were determined based on the presence of the peak in at least 5% of the cells for that cell type. The total number of peaks in each gene plus promoter region was quantified per gene for each cell type and compared to the number of peaks in the merged peak set for each gene set using a Fisher’s exact test. The peaks in the remaining gene plus promoter regions were used as background. Gene sets from Coe et al.^22^ (COE253), Kaplanis et al.^23^ (DDD299) and SFARI gene (SFARI854) were used for enrichment testing. P-values were Bonferroni corrected for 81 tests (27 cell types and 3 gene sets). In addition, we also compared the number of probands and siblings carrying one or more DNMs in a peak within the gene plus promoter region of genes in the union set of the three gene sets tested above. Burden was quantified using a Fisher’s exact test for each peak set. P-values were Bonferroni corrected for 28 tests (# of cell types plus the merged peak set).

#### Morbidity map CNV enrichment

CNVs enriched in NDD cases from Coe et al 2014^22^ were intersected with peak sets using bedtools 2.24.0; peaks were required to have a 50% overlap with the CNV region. Peak sets were determined based on the presence of the peak in at least 5% of the cells for that cell type. The total number of peaks overlapping a CNV were compared to the number of peaks that did not overlap with a CNV for each cell type. The merged peak set was used as background and compared by Fisher’s exact test. P-values were Bonferroni corrected for 27 tests (# of cell types).

#### Cell type-specific GWAS enrichment testing

We retrieved GWAS summary statistics for schizophrenia (Ripke et al., 2014)^24^, bipolar disorder (Stahl et al., 2019)^25^, and autism (Grove et al., 2019)^26^ from the Psychiatric Genomics Consortium data portal (https://www.med.unc.edu/pgc). We also obtained GWAS summary statistics for schizophrenia (Pardiñas et al., 2018)^27^ from http://walters.psycm.cf.ac.uk/. GWAS summary statistics for major depression (Howard et al., 2019)^28^ were obtained from the authors under the auspices of a Data Use Agreement between 23AndMe and the University of Maryland Baltimore. We applied stratified LD score regression (LDSC version 1.0.1; Finucane et al., 2018; Finucane et al., 2015)^29, 30^ to these summary statistics to evaluate the enrichment of trait heritability in each of 27 cell type-specific peak sets, which we defined as peaks present in at least 1% of cells from a given cell type. These associations were adjusted for the union of the peak sets as well as for 52 annotations from version 1.2 of the LDSC baseline model (including genic regions, enhancer regions and conserved regions; Finucane et al., 2015)^29^. Associations that met a cutoff of FDR < 0.05 were considered significant.

## EXTENDED DATA FIGURES

**Extended Data Fig. 1:**
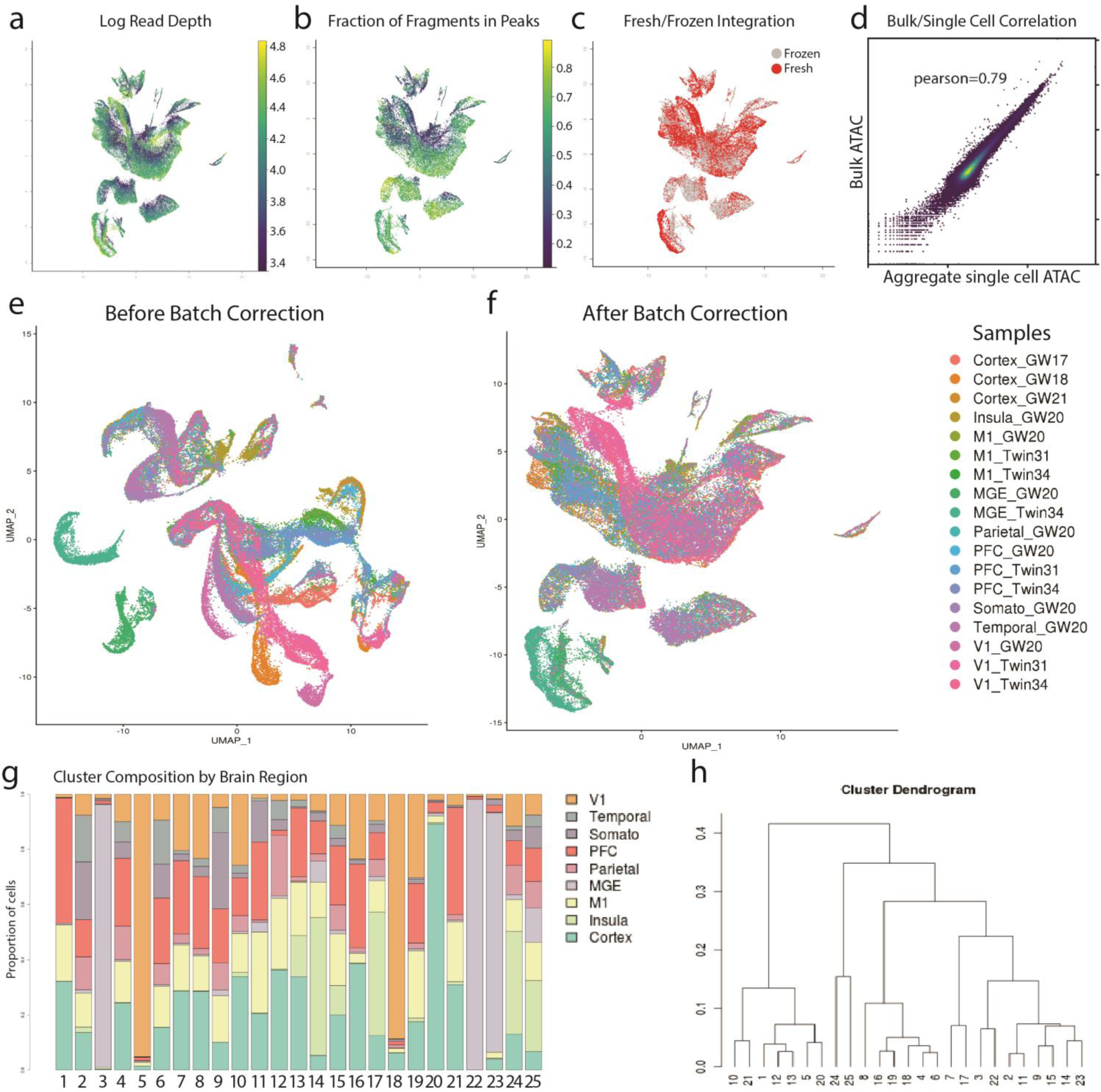
Batch correction and quality control metrics for primary scATACseq data. **a)** UMAP projection of all primary scATACseq cells colored by log10(read depth). **b)** UMAP projection of fractions of reads in peaks for all primary scATACseq cells. **c)** UMAP projection of all primary scATACseq cells colored by condition (fresh/frozen). **d)** Aggregate signal of single cell data was highly correlated with bulk ATACseq libraries prepared in parallel. Pearson’s correlation coefficient (*r*=0.79) between bulk ATAC-seq and aggregate of all scATACseq cells was calculated from the PFC_GW20 sample based on coverage of 10Kb genomic bins. Bulk ATAC-seq and scATACseq data were generated from the same sample. **e)** UMAP projection of all primary scATACseq cells before batch correction colored by sample. **f)** UMAP projection of all primary scATACseq cells after batch correction colored by sample (Methods). **g)** Barplot depicting the proportion of cells from each brain region for each leiden cluster for all primary scATACseq cells. **h)** Dendrogram of hierarchical clustering of leiden clusters for all primary scATACseq cells.

**Extended Data Fig. 2:**
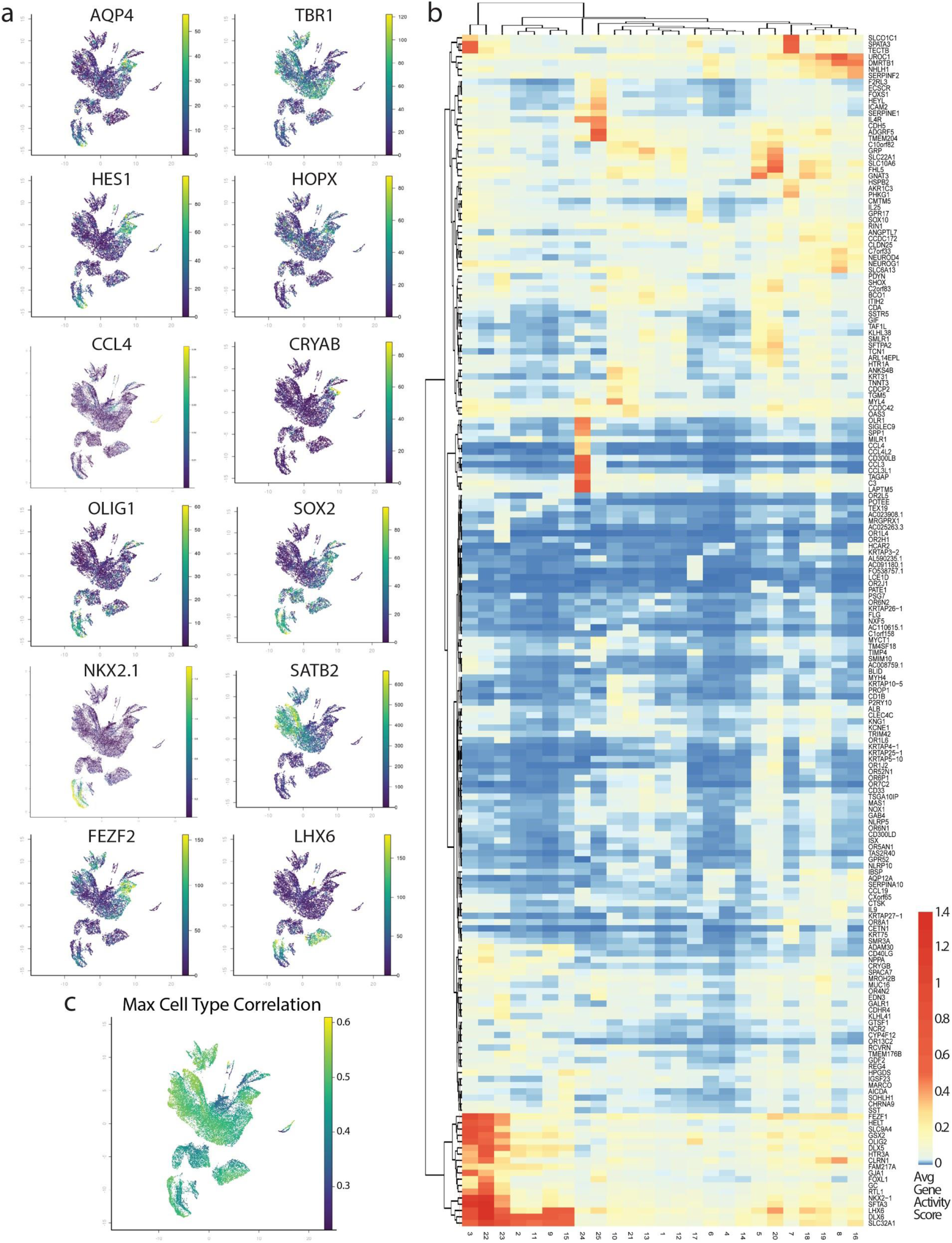
Gene activity scores correlate with cell type-specific expression of marker genes. **a)** UMAP projections of all primary scATACseq cells colored by gene activity score. From top left to bottom right, AQP4 marking glia/astrocytes, TBR1 marking excitatory neurons, HES1 marking radial glia, HOPX marking outer radial glia, CCL4 marking microglia, CRYAB marking truncated radial glia, OLIG1 marking oligodendrocyte precursors, SOX2 marking radial glia, NKX2.1 marking MGE cells, SATB2 marking upper layer excitatory neurons, FEZF2 marking deep layer excitatory neurons, and LHX6 marking MGE-derived interneurons. **b)** Heatmap of average gene activity scores for 200 variable genes for the 25 leiden clusters of all primary scATACseq cells. **c)** UMAP projection of max correlation values with scRNAseq cell types.

**Extended Data Fig. 3:**
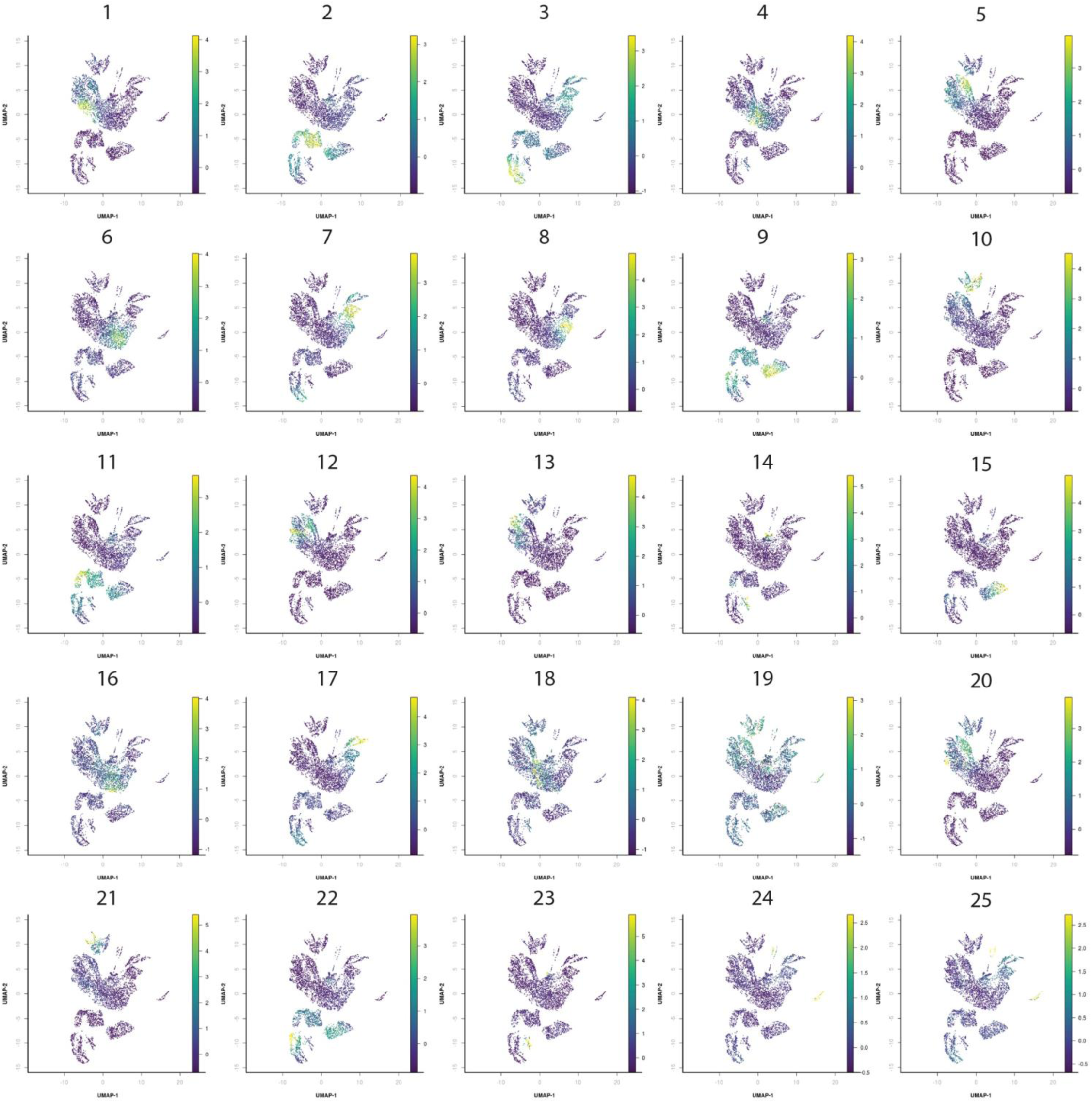
Projection of cluster specific peaks for all scATACseq clusters. UMAP projections of all primary scATACseq cells colored by Z-score of peak set enrichment. From top left to bottom right, projection of cluster specific peak sets for each of the 25 leiden clusters (Fisher’s Exact, two-sided, FDR<0.05).

**Extended Data Fig. 4:**
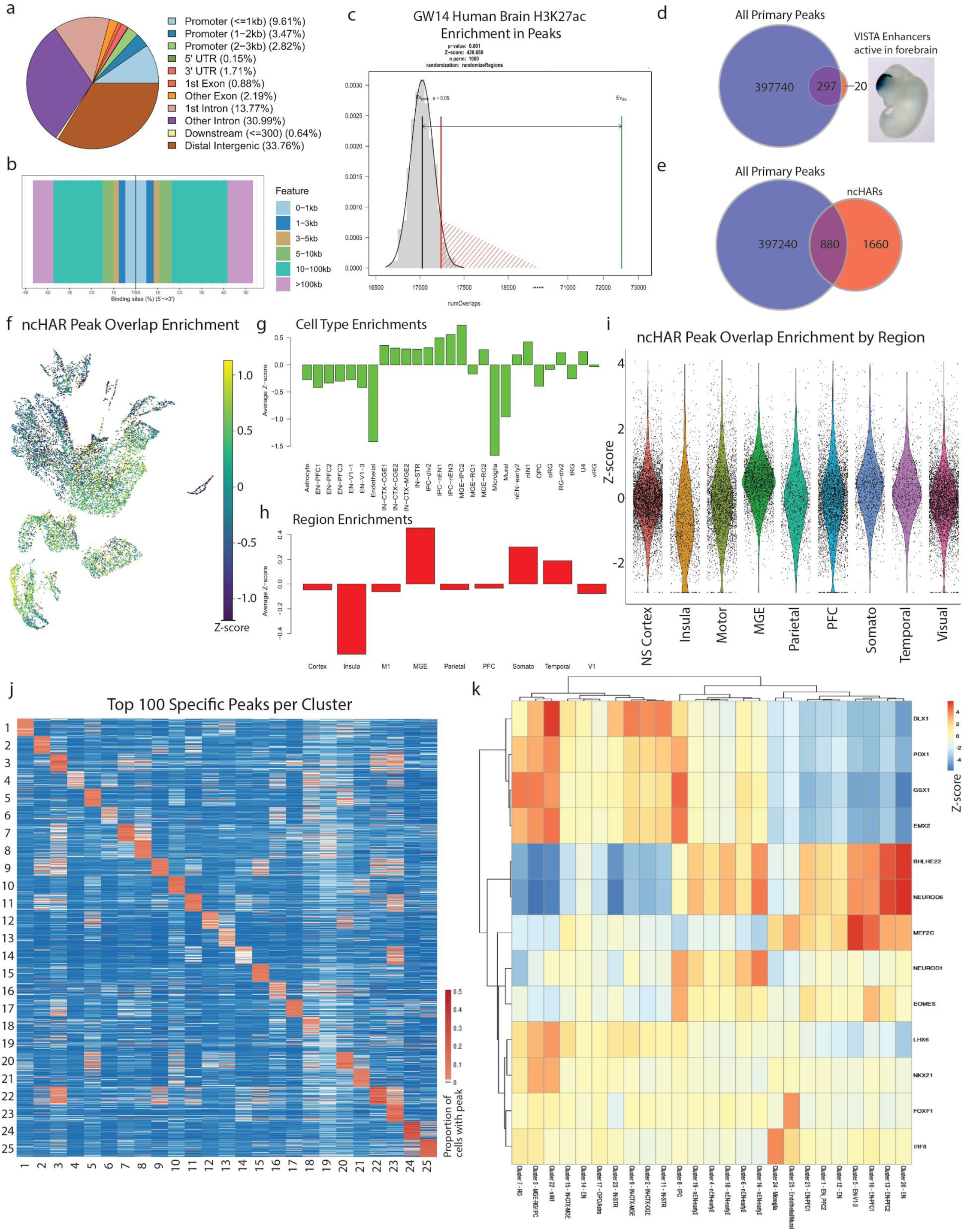
scATACseq peaks overlap with marks of active enhancers and previously validated forebrain enhancers. **a)** Annotation of peaks called from all primary scATACseq cells in genomic features. **b)** Distribution of peaks called from all primary scATACseq cells in flanking regions around transcription start sites. **c)** Enrichment of overlaps of H3K27ac ChIP-seq peaks generated from GW14 human frontal cortex (GEO: GSE63648) with peaks called from all primary scATACseq cells (Permutation Test, one-sided, p<0.001). **d)** Overlap of 317 VISTA enhancer regions active in the forebrain with peaks called from all primary scATACseq cells (297/217, Image taken from VISTA Enhancer Browser, hs123 - embryo 1). **e)** Overlap of 2540 non-coding human accelerated regions (ncHARS) taken from Capra et al. 2013 with peaks called from all primary scATACseq cells (880/2540). **f)** UMAP projection of all primary scATACseq cells colored by Z-score of enrichment of peaks overlapping ncHARs. **g)** Average Z-score of enrichment of peaks overlapping ncHARs grouped by cell type predictions. h) Average Z-score of enrichment of peaks overlapping ncHARs grouped by brain region. **i)** Violin plots depicting Z-scores of enrichment of peaks overlapping ncHARs grouped by brain region. **j)** Heatmap of average proportion of cells in each cluster that have reads overlapping top cluster specific peaks (Fisher’s Exact). Columns represent the signal from each cluster arranged left-to-right 1-25. Rows represent top 100 cluster specific peaks for each cluster arranged top-to-bottom 1-25. **k)** Average transcription factor motif enrichments for 13 TFs in 25 leiden clusters from all primary scATACseq cells calculated using ChromVAR (Methods).

**Extended Data Figure 5:**
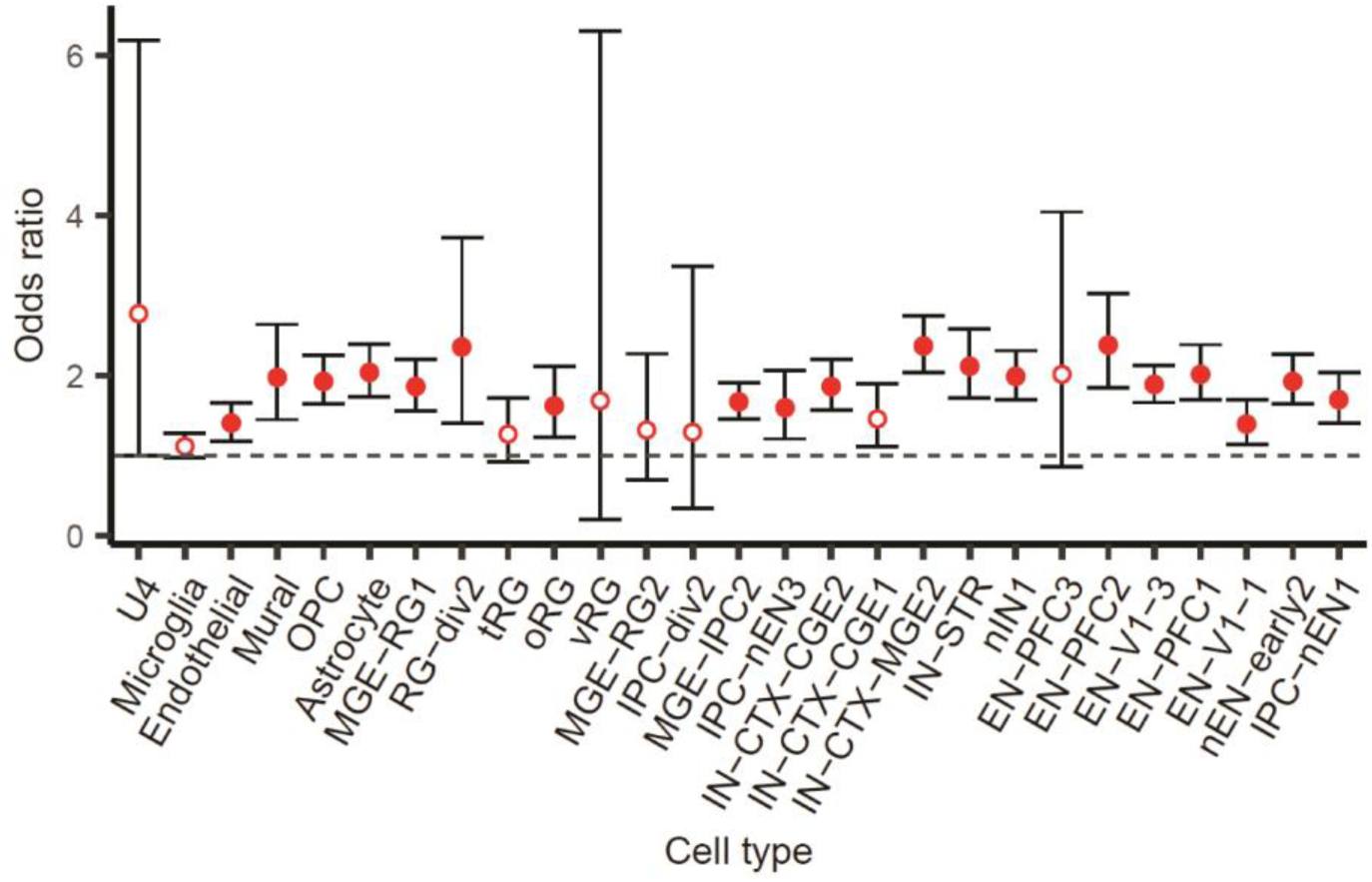
DNM enrichment in cell type-specific peaks. Peaks in 19 out of 27 cell types are enriched for DNMs observed in ASD cases as compared to the merged peak set. Filled circles indicate cell types with significant enrichment after Bonferroni correction. We did not find any differences in the number of DNMs in cell type-specific peaks between probands and siblings across any cell type after Bonferroni correction.

**Extended Data Fig. 6:**
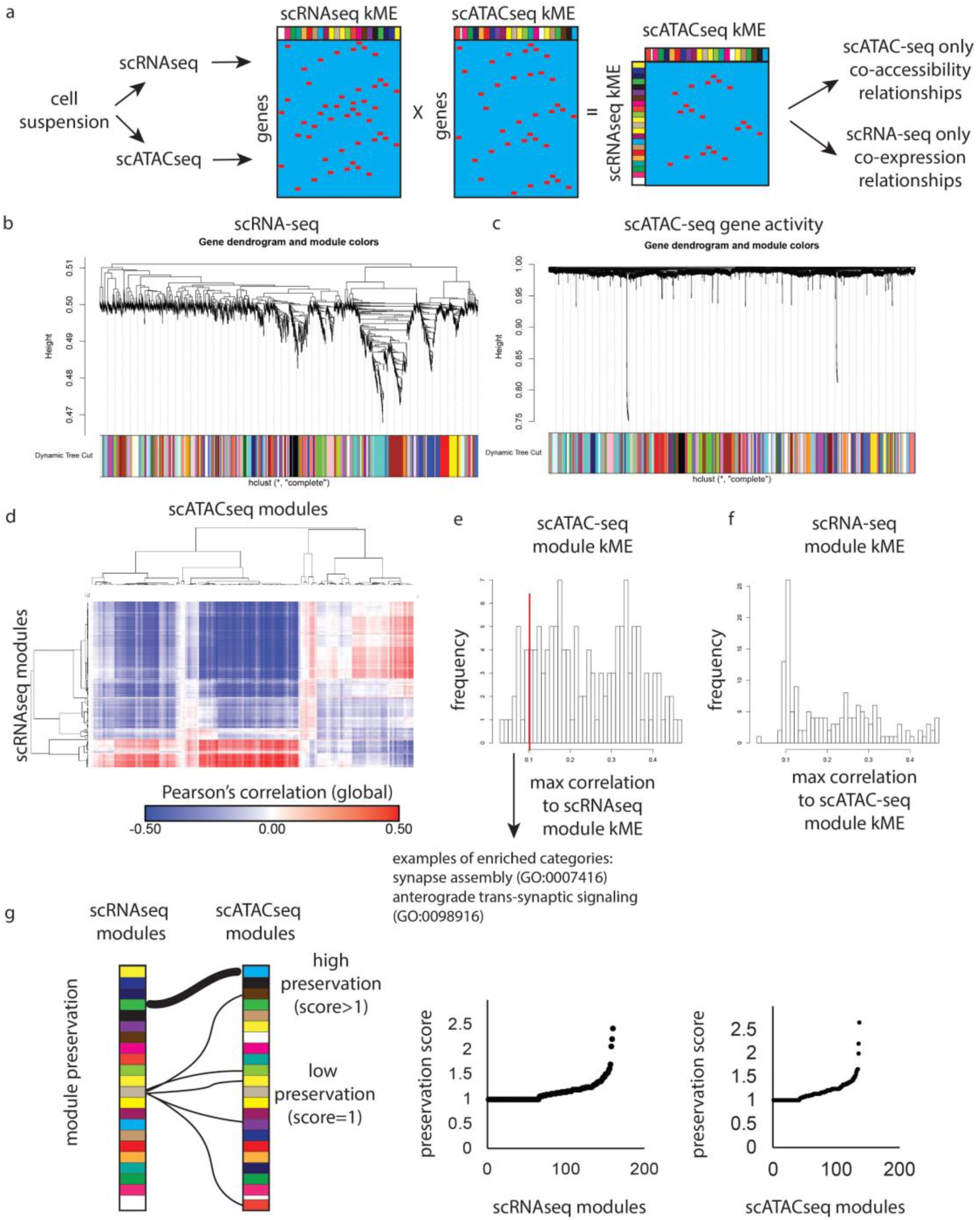
Preservation of gene co-expression relationships inferred from transcriptomic and chromatin state profiling of single cells. **a)** Single cells profiled for scRNAseq and scATACseq were analyzed for gene co-expression relationships using weighted gene coexpression network analysis^31^. For scATACseq, we used gene activity scores as proxy for mRNA expression. **b-c)** WGCNA hierarchical clustering plots and module assignments of genes **(b)** or gene activity scores **(c)**. See also Supplementary Tables 3 and 4 for gene-module assignments and module eigengene correlations. **d)** Gene-module correlations were compared between scRNAseq and scATACseq datasets, revealing a high degree of correlation between gene co-expression modules calculated from scRNAseq and those inferred from gene activity scores in scATACseq data. **e-f)** Histograms showing the distributions of module correlations across modalities. In either comparison, only a small number of clusters lack appreciable correlation to any module in the other modality, with scATACseq-specific modules enriched for genes involved in cell-cell interactions, including the protocadherin gene clusters that are highly co-accessible across single cells, but not highly co-expressed. **g)** Analysis of gene-module memberships reveal high degree of conserved gene co-expression and co-accessibility across single cells.

**Extended Data Fig. 7:**
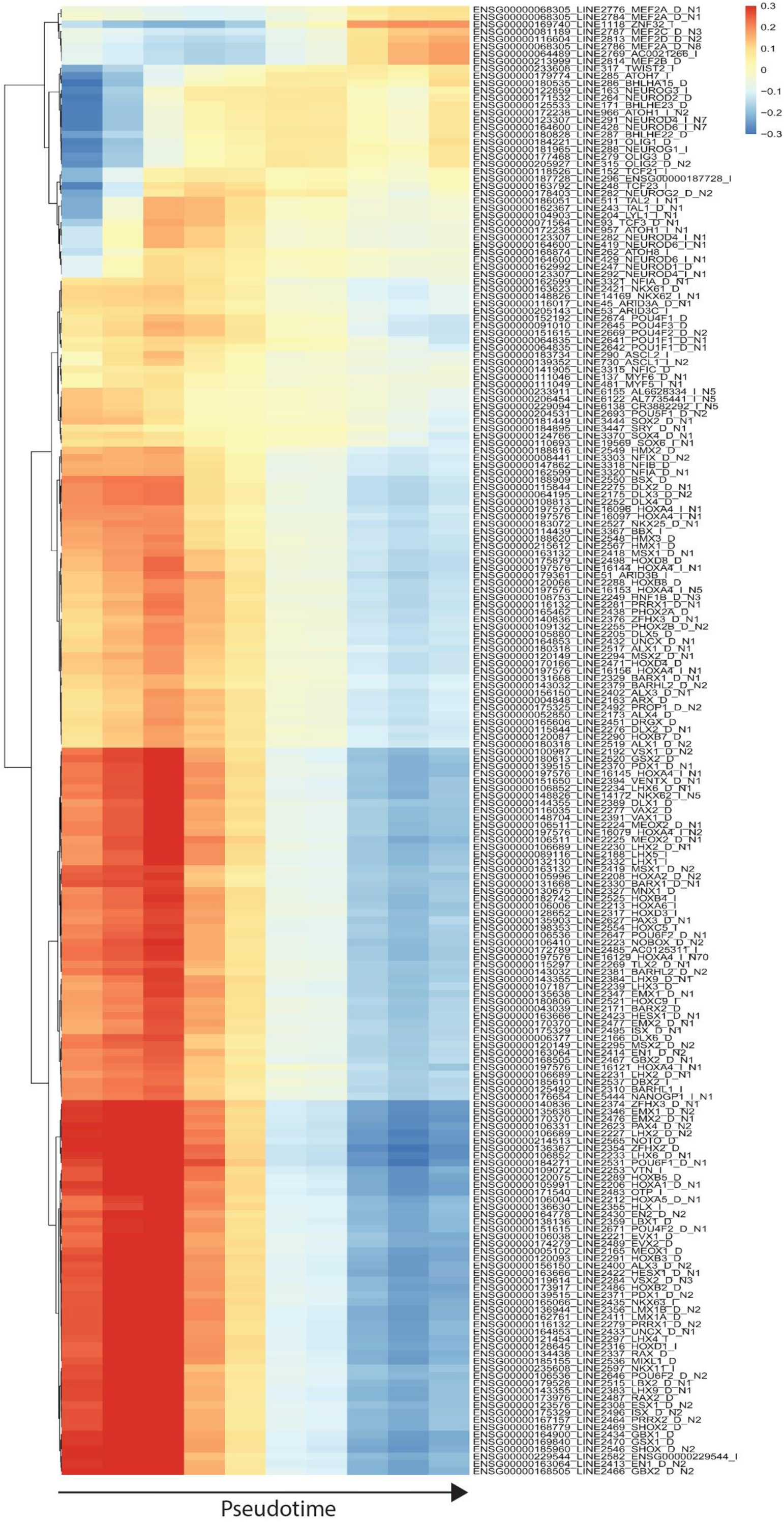
Temporally dynamic transcription factor motif enrichments reveal regulators of excitatory differentiation. Heatmap of average transcription factor motif enrichments for the top 200 most variable transcription factors as determined by ChromVAR (Methods) in equally-sized bins of pseudotime arranged in increasing order from left-to-right.

**Extended Data Fig. 8:**
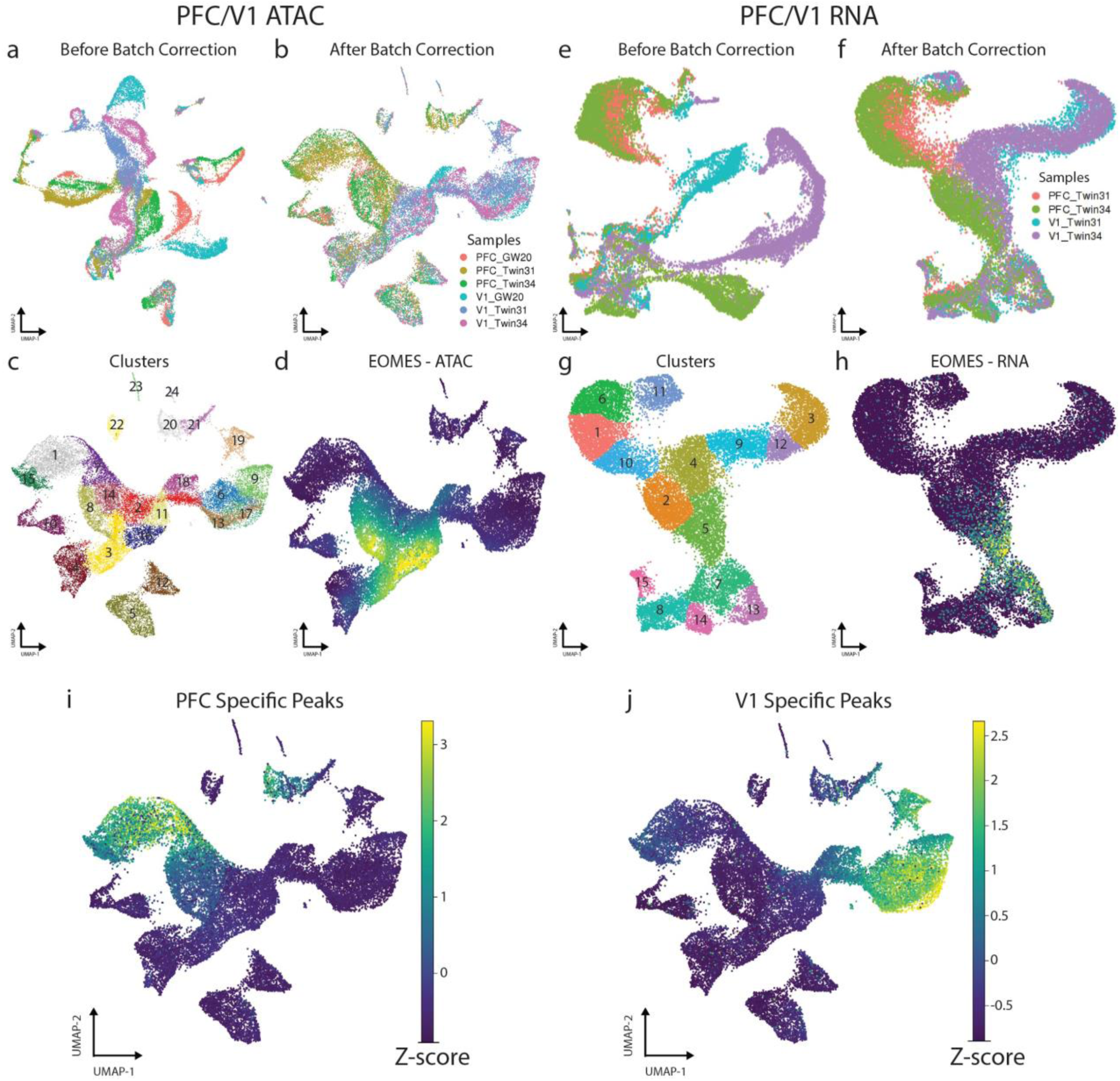
Chromatin state profiling reveals divergence of PFC and V1 excitatory lineages. **a)** UMAP projection of scATACseq cells from PFC and V1 samples before batch correction colored by sample. **b)** UMAP projection of scATACseq cells from PFC and V1 samples after batch correction colored by sample. **c)** UMAP projection of scATACseq cells from PFC and V1 samples colored by leiden cluster. **d)** UMAP projection of scATACseq cells from PFC and V1 samples colored by gene activity score for EOMES, a marker of IPCs. **e)** UMAP projection of scRNAseq cells from PFC and V1 samples before batch correction colored by sample. **f)** UMAP projection of scRNAseq cells from PFC and V1 samples after batch correction colored by sample. **g)** UMAP projection of scRNAseq cells from PFC and V1 samples colored by leiden cluster. **h)** UMAP projection of scRNAseq cells from PFC and V1 samples colored by expression of EOMES, a marker of IPCs. **i)** UMAP projection of scATACseq cells from PFC and V1 samples colored by Z-score of enrichment of 5,863 PFC-specific peaks (Fisher’s Exact, two-sided, FDR<0.05). **j)** UMAP projection of scATACseq cells from PFC and V1 samples colored by Z-score of enrichment of 26,520 V1-specific peaks (Fisher’s Exact, two-sided, FDR<0.05)

**Extended Data Figure 9:**
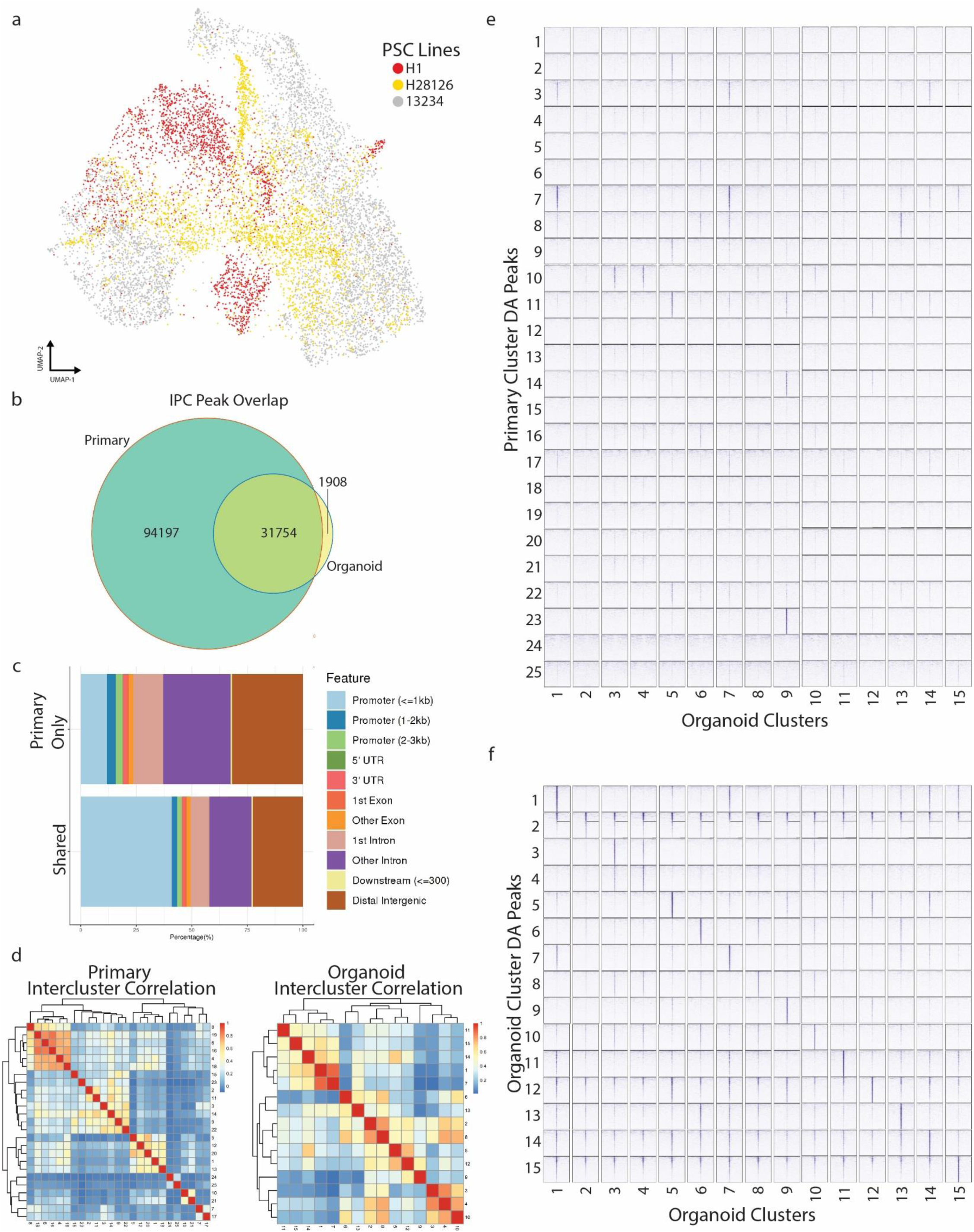
Comparison of organoid and primary peaks reveal significant differences in the chromatin landscapes. **a)** UMAP projection of all organoid cells colored by sample of origin. PSC – pluripotent stem cell. **b)** Venn diagram depicting the overlap of peaks called from IPCs from primary samples with peaks called from IPCs from organoid samples. **c)** Annotation of peaks found only in primary IPCs (top) and peaks shared between primary and organoids (bottom) in genomic features. **d)** Left, heatmap of correlations between leiden clusters of all primary scATACseq cells. Right, heatmap of correlations between leiden clusters of all organoid scATACseq cells. The average inter-cluster correlation for primary clusters (*r*=0.2) is approximately half that of organoid clusters (*r*=0.39). **e)** Pileups of ATACseq signal from each organoid leiden cluster in sets of top 1000 DA peaks for each primary leiden cluster. Pileups are centered on each peak and the flanking +/-10Kb regions are shown. **f)** Pileups of ATACseq signal from each organoid leiden cluster in sets of top 1000 DA peaks for each organoid leiden cluster. Pileups are centered on each peak and the flanking +/-10Kb regions are shown.

**Extended Data Table 1:**
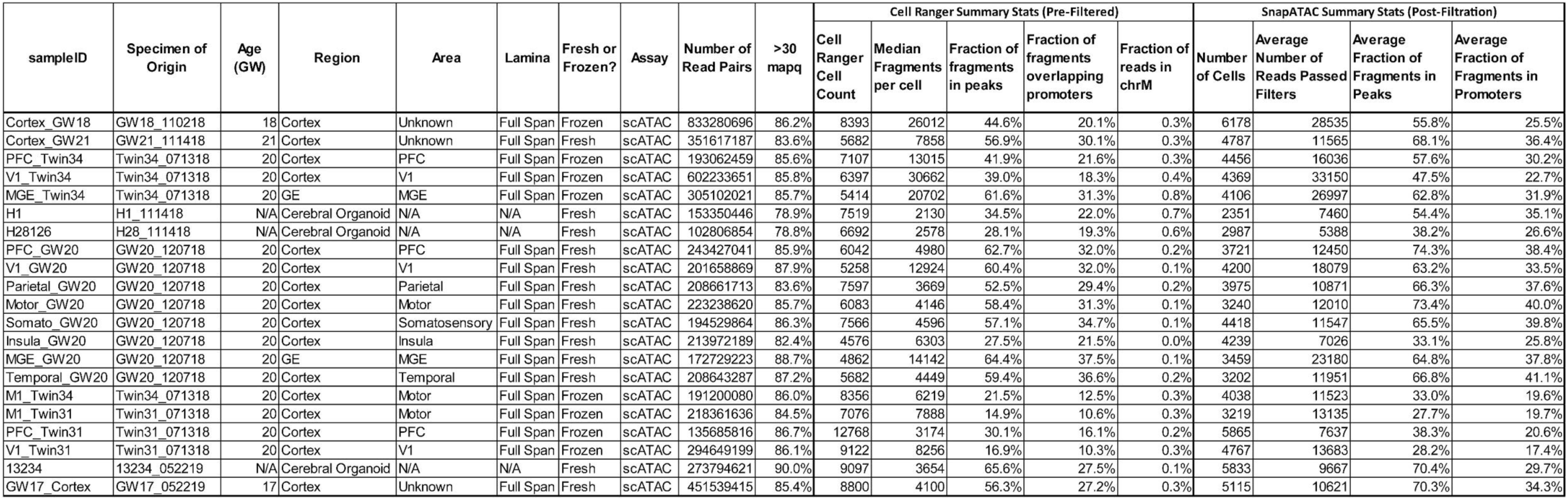
Sample Metadata Table. Includes sample metadata and key summary statistics including sequencing depth, cell counts, fraction of fragments in peaks, and fraction of fragments in promoters.

## References

1. Nowakowski, T. J. et al. Spatiotemporal gene expression trajectories reveal developmental hierarchies of the human cortex. Science 358, 1318–1323, doi:10.1126/science.aap8809 (2017).

2. Tasic, B. et al. Shared and distinct transcriptomic cell types across neocortical areas. Nature 563, 72–78, doi:10.1038/s41586-018-0654-5 (2018).

3. Hodge, R. D. et al. Conserved cell types with divergent features in human versus mouse cortex. Nature 573, 61–68, doi:10.1038/s41586-019-1506-7 (2019).

4. Thomsen, E. R. et al. Fixed single-cell transcriptomic characterization of human radial glial diversity. Nat Methods 13, 87–93, doi:10.1038/nmeth.3629 (2016).

5. Nowakowski, T. J., Pollen, A. A., Sandoval-Espinosa, C. & Kriegstein, A. R. Transformation of the Radial Glia Scaffold Demarcates Two Stages of Human Cerebral Cortex Development. Neuron 91, 1219–1227, doi:10.1016/j.neuron.2016.09.005 (2016).

6. Pollen, A. A. et al. Molecular identity of human outer radial glia during cortical development. Cell 163, 55–67, doi:10.1016/j.cell.2015.09.004 (2015).

7. Waddington, C. H. The strategy of the genes. (Routledge, 2014).

8. Visel, A. et al. A high-resolution enhancer atlas of the developing telencephalon. Cell 152, 895–908, doi:10.1016/j.cell.2012.12.041 (2013).

9. Pattabiraman, K. et al. Transcriptional regulation of enhancers active in protodomains of the developing cerebral cortex. Neuron 82, 989–1003, doi:10.1016/j.neuron.2014.04.014 (2014).

10. Buenrostro, J. D., Giresi, P. G., Zaba, L. C., Chang, H. Y. & Greenleaf, W. J. Transposition of native chromatin for fast and sensitive epigenomic profiling of open chromatin, DNA-binding proteins and nucleosome position. Nat Methods 10, 1213–1218, doi:10.1038/nmeth.2688 (2013).

11. Johnson, D. S., Mortazavi, A., Myers, R. M. & Wold, B. Genome-wide mapping of in vivo protein-DNA interactions. Science 316, 1497–1502, doi:10.1126/science.1141319 (2007).

12. Bonev, B. et al. Multiscale 3D Genome Rewiring during Mouse Neural Development. Cell 171, 557–572 e524, doi:10.1016/j.cell.2017.09.043 (2017).

13. Mo, A. et al. Epigenomic Signatures of Neuronal Diversity in the Mammalian Brain. Neuron 86, 1369–1384, doi:10.1016/j.neuron.2015.05.018 (2015).

14. de la Torre-Ubieta, L. et al. The Dynamic Landscape of Open Chromatin during Human Cortical Neurogenesis. Cell 172, 289–304 e218, doi:10.1016/j.cell.2017.12.014 (2018).

15. Luo, C. et al. Cerebral Organoids Recapitulate Epigenomic Signatures of the Human Fetal Brain. Cell Rep 17, 3369–3384, doi:10.1016/j.celrep.2016.12.001 (2016).

16. Nord, A. S. et al. Rapid and pervasive changes in genome-wide enhancer usage during mammalian development. Cell 155, 1521–1531, doi:10.1016/j.cell.2013.11.033 (2013).

17. Amiri, A. et al. Transcriptome and epigenome landscape of human cortical development modeled in organoids. Science 362, doi:10.1126/science.aat6720 (2018).

18. Buenrostro, J. D. et al. Single-cell chromatin accessibility reveals principles of regulatory variation. Nature 523, 486–490, doi:10.1038/nature14590 (2015).

19. Cusanovich, D. A. et al. Multiplex single cell profiling of chromatin accessibility by combinatorial cellular indexing. Science 348, 910–914, doi:10.1126/science.aab1601 (2015).

20. Preissl, S. et al. Single-nucleus analysis of accessible chromatin in developing mouse forebrain reveals cell-type-specific transcriptional regulation. Nat Neurosci 21, 432–439, doi:10.1038/s41593-018-0079-3 (2018).

21. Zhu, C. et al. An ultra high-throughput method for single-cell joint analysis of open chromatin and transcriptome. Nat Struct Mol Biol 26, 1063–1070, doi:10.1038/s41594-019-0323-x (2019).

22. Lake, B. B. et al. Integrative single-cell analysis of transcriptional and epigenetic states in the human adult brain. Nat Biotechnol 36, 70–80, doi:10.1038/nbt.4038 (2018).

23. Pollard, K. S. et al. Forces shaping the fastest evolving regions in the human genome. PLoS Genet 2, e168, doi:10.1371/journal.pgen.0020168 (2006).

24. Capra, J. A., Erwin, G. D., McKinsey, G., Rubenstein, J. L. & Pollard, K. S. Many human accelerated regions are developmental enhancers. Philos Trans R Soc Lond B Biol Sci 368, 20130025, doi:10.1098/rstb.2013.0025 (2013).

25. Reilly, S. K., et al. Evolutionary genomics. Evolutionary changes in promoter and enhancer activity during human corticogenesis. Science 347, 1155–1159, doi:10.1126/science.1260943 (2015).

26. Doan, R. N. et al. Mutations in Human Accelerated Regions Disrupt Cognition and Social Behavior. Cell 167, 341–354 e312, doi:10.1016/j.cell.2016.08.071 (2016).

27. Hansen, D. V. et al. Non-epithelial stem cells and cortical interneuron production in the human ganglionic eminences. Nat Neurosci 16, 1576–1587, doi:10.1038/nn.3541 (2013).

28. Ma, T. et al. Subcortical origins of human and monkey neocortical interneurons. Nat Neurosci 16, 1588–1597, doi:10.1038/nn.3536 (2013).

29. Johansen, N. & Quon, G. scAlign: a tool for alignment, integration, and rare cell identification from scRNA-seq data. Genome Biol 20, 166, doi:10.1186/s13059-019-1766-4 (2019).

30. Sinnamon, J. R. et al. The accessible chromatin landscape of the murine hippocampus at single-cell resolution. Genome Res 29, 857–869, doi:10.1101/gr.243725.118 (2019).

31. Zhang, Y. et al. Model-based analysis of ChIP-Seq (MACS). Genome Biol 9, R137, doi:10.1186/gb-2008-9-9-r137 (2008).

32. Visel, A., Minovitsky, S., Dubchak, I. & Pennacchio, L. A. VISTA Enhancer Browser--a database of tissue-specific human enhancers. Nucleic Acids Res 35, D88–92, doi:10.1093/nar/gkl822 (2007).

33. Sousa, A. M. M. et al. Molecular and cellular reorganization of neural circuits in the human lineage. Science 358, 1027–1032, doi:10.1126/science.aan3456 (2017).

34. Krienen, F. M. et al. Innovations in Primate Interneuron Repertoire. bioRxiv, 709501, doi:10.1101/709501 (2019).

35. Heinz, S. et al. Simple combinations of lineage-determining transcription factors prime cis-regulatory elements required for macrophage and B cell identities. Mol Cell 38, 576–589, doi:10.1016/j.molcel.2010.05.004 (2010).

36. Fulco, C. P. et al. Activity-by-contact model of enhancer-promoter regulation from thousands of CRISPR perturbations. Nat Genet 51, 1664–1669, doi:10.1038/s41588-019-0538-0 (2019).

37. Schizophrenia Working Group of the Psychiatric Genomics, C. Biological insights from 108 schizophrenia-associated genetic loci. Nature 511, 421–427, doi:10.1038/nature13595 (2014).

38. Satterstrom, F. K. et al. Large-scale exome sequencing study implicates both developmental and functional changes in the neurobiology of autism. bioRxiv, 484113, doi:10.1101/484113 (2019).

39. Turner, T. N. et al. Sex-Based Analysis of De Novo Variants in Neurodevelopmental Disorders. Am J Hum Genet 105, 1274–1285, doi:10.1016/j.ajhg.2019.11.003 (2019).

40. Werling, D. M. et al. An analytical framework for whole-genome sequence association studies and its implications for autism spectrum disorder. Nat Genet 50, 727–736, doi:10.1038/s41588-018-0107-y (2018).

41. Stessman, H. A. et al. Targeted sequencing identifies 91 neurodevelopmental-disorder risk genes with autism and developmental-disability biases. Nat Genet 49, 515–526, doi:10.1038/ng.3792 (2017).

42. Coe, B. P. et al. Refining analyses of copy number variation identifies specific genes associated with developmental delay. Nat Genet 46, 1063–1071, doi:10.1038/ng.3092 (2014).

43. Pollen, A. A. et al. Establishing Cerebral Organoids as Models of Human-Specific Brain Evolution. Cell 176, 743–756 e717, doi:10.1016/j.cell.2019.01.017 (2019).

44. Langfelder, P. & Horvath, S. WGCNA: an R package for weighted correlation network analysis. BMC Bioinformatics 9, 559, doi:10.1186/1471-2105-9-559 (2008).

45. Cao, J. et al. The single-cell transcriptional landscape of mammalian organogenesis. Nature 566, 496–502, doi:10.1038/s41586-019-0969-x (2019).

46. Nord, A. S. & West, A. E. Neurobiological functions of transcriptional enhancers. Nat Neurosci 23, 5–14, doi:10.1038/s41593-019-0538-5 (2020).

47. Inoue, F., Kreimer, A., Ashuach, T., Ahituv, N. & Yosef, N. Identification and Massively Parallel Characterization of Regulatory Elements Driving Neural Induction. Cell Stem Cell 25, 713–727 e710, doi:10.1016/j.stem.2019.09.010 (2019).

48. Miller, J. A. et al. Transcriptional landscape of the prenatal human brain. Nature 508, 199–206, doi:10.1038/nature13185 (2014).

49. Morriss, G. M. Morphogenesis of the malformations induced in rat embryos by maternal hypervitaminosis A. J Anat 113, 241–250 (1972).

50. Osei-Sarfo, K. & Gudas, L. J. Retinoic acid suppresses the canonical Wnt signaling pathway in embryonic stem cells and activates the noncanonical Wnt signaling pathway. Stem Cells 32, 2061–2071, doi:10.1002/stem.1706 (2014).

51. Harrison-Uy, S. J., Siegenthaler, J. A., Faedo, A., Rubenstein, J. L. & Pleasure, S. J. CoupTFI interacts with retinoic acid signaling during cortical development. PLoS One 8, e58219, doi:10.1371/journal.pone.0058219 (2013).

52. Bartholin, L. et al. TGIF inhibits retinoid signaling. Mol Cell Biol 26, 990–1001, doi:10.1128/MCB.26.3.990-1001.2006 (2006).

53. Camp, J. G. et al. Human cerebral organoids recapitulate gene expression programs of fetal neocortex development. Proc Natl Acad Sci U S A 112, 15672–15677, doi:10.1073/pnas.1520760112 (2015).

54. Kanton, S. et al. Organoid single-cell genomic atlas uncovers human-specific features of brain development. Nature 574, 418–422, doi:10.1038/s41586-019-1654-9 (2019).

55. Skene, N. G. et al. Genetic identification of brain cell types underlying schizophrenia. Nat Genet 50, 825–833, doi:10.1038/s41588-018-0129-5 (2018).

56. Li, M. et al. Integrative functional genomic analysis of human brain development and neuropsychiatric risks. Science 362, doi:10.1126/science.aat7615 (2018).

57. Hrvatin, S. et al. PESCA: A scalable platform for the development of cell-type-specific viral drivers. bioRxiv, 570895, doi:10.1101/570895 (2019).

58. Mich, J. K. et al. Epigenetic landscape and AAV targeting of human neocortical cell classes. bioRxiv, 555318, doi:10.1101/555318 (2019).

59. Graybuck, L. T. et al. Prospective, brain-wide labeling of neuronal subclasses with enhancer-driven AAVs. bioRxiv, 525014, doi:10.1101/525014 (2019).

60. Kaplanis, J. et al. Integrating healthcare and research genetic data empowers the discovery of 49 novel developmental disorders. bioRxiv, 797787, doi:10.1101/797787 (2019).

61. Coe, B. P. et al. Neurodevelopmental disease genes implicated by de novo mutation and copy number variation morbidity. Nat Genet 51, 106–116, doi:10.1038/s41588-018-0288-4 (2019).

62. Howard, D. M. et al. Genome-wide meta-analysis of depression identifies 102 independent variants and highlights the importance of the prefrontal brain regions. Nat Neurosci 22, 343–352, doi:10.1038/s41593-018-0326-7 (2019).

63. Pardinas, A. F. et al. Common schizophrenia alleles are enriched in mutation-intolerant genes and in regions under strong background selection. Nat Genet 50, 381–389, doi:10.1038/s41588-018-0059-2 (2018).

64. Stahl, E. A. et al. Genome-wide association study identifies 30 loci associated with bipolar disorder. Nat Genet 51, 793–803, doi:10.1038/s41588-019-0397-8 (2019).

## References

1. Corces, M. R. et al. An improved ATAC-seq protocol reduces background and enables interrogation of frozen tissues. Nat Methods 14, 959–962, doi:10.1038/nmeth.4396 (2017).

2. Kadoshima, T. et al. Self-organization of axial polarity, inside-out layer pattern, and species-specific progenitor dynamics in human ES cell-derived neocortex. Proc Natl Acad Sci U S A 110, 20284–20289, doi:10.1073/pnas.1315710110 (2013).

3. Pollen, A. A. et al. Establishing Cerebral Organoids as Models of Human-Specific Brain Evolution. Cell 176, 743–756 e717, doi:10.1016/j.cell.2019.01.017 (2019).

4. Wolock, S. L., Lopez, R. & Klein, A. M. Scrublet: Computational Identification of Cell Doublets in Single-Cell Transcriptomic Data. Cell Syst 8, 281–291 e289, doi:10.1016/j.cels.2018.11.005 (2019).

5. Hafemeister, C. & Satija, R. Normalization and variance stabilization of single-cell RNA-seq data using regularized negative binomial regression. bioRxiv, 576827, doi:10.1101/576827 (2019).

6. Butler, A., Hoffman, P., Smibert, P., Papalexi, E. & Satija, R. Integrating single-cell transcriptomic data across different conditions, technologies, and species. Nat Biotechnol 36, 411–420, doi:10.1038/nbt.4096 (2018).

7. Fang, R. et al. Fast and Accurate Clustering of Single Cell Epigenomes Reveals Cis-Regulatory Elements in Rare Cell Types. bioRxiv, 615179, doi:10.1101/615179 (2019).

8. Johansen, N. & Quon, G. scAlign: a tool for alignment, integration, and rare cell identification from scRNA-seq data. Genome Biol 20, 166, doi:10.1186/s13059-019-1766-4 (2019).

9. van Dijk, D. et al. Recovering Gene Interactions from Single-Cell Data Using Data Diffusion. Cell 174, 716–729 e727, doi:10.1016/j.cell.2018.05.061 (2018).

10. Nowakowski, T. J. et al. Spatiotemporal gene expression trajectories reveal developmental hierarchies of the human cortex. Science 358, 1318–1323, doi:10.1126/science.aap8809 (2017).

11. Ramirez, F. et al. deepTools2: a next generation web server for deep-sequencing data analysis. Nucleic Acids Res 44, W160–165, doi:10.1093/nar/gkw257 (2016).

12. Heinz, S. et al. Simple combinations of lineage-determining transcription factors prime cis-regulatory elements required for macrophage and B cell identities. Mol Cell 38, 576–589, doi:10.1016/j.molcel.2010.05.004 (2010).

13. Schep, A. N., Wu, B., Buenrostro, J. D. & Greenleaf, W. J. chromVAR: inferring transcription-factor-associated accessibility from single-cell epigenomic data. Nat Methods 14, 975–978, doi:10.1038/nmeth.4401 (2017).

14. Fulco, C. P. et al. Activity-by-contact model of enhancer-promoter regulation from thousands of CRISPR perturbations. Nat Genet 51, 1664–1669, doi:10.1038/s41588-019-0538-0 (2019).

15. Capra, J. A., Erwin, G. D., McKinsey, G., Rubenstein, J. L. & Pollard, K. S. Many human accelerated regions are developmental enhancers. Philos Trans R Soc Lond B Biol Sci 368, 20130025, doi:10.1098/rstb.2013.0025 (2013).

16. Reilly, S. K., et al. Evolutionary genomics. Evolutionary changes in promoter and enhancer activity during human corticogenesis. Science 347, 1155–1159, doi:10.1126/science.1260943 (2015).

17. Gel, B. et al. regioneR: an R/Bioconductor package for the association analysis of genomic regions based on permutation tests. Bioinformatics 32, 289–291, doi:10.1093/bioinformatics/btv562 (2016).

18. Yu, G., Wang, L. G. & He, Q. Y. ChIPseeker: an R/Bioconductor package for ChIP peak annotation, comparison and visualization. Bioinformatics 31, 2382–2383, doi:10.1093/bioinformatics/btv145 (2015).

19. Cao, J. et al. The single-cell transcriptional landscape of mammalian organogenesis. Nature 566, 496–502, doi:10.1038/s41586-019-0969-x (2019).

20. Pliner, H. A. et al. Cicero Predicts cis-Regulatory DNA Interactions from Single-Cell Chromatin Accessibility Data. Mol Cell 71, 858–871 e858, doi:10.1016/j.molcel.2018.06.044 (2018).

21. Farrell, J. A. et al. Single-cell reconstruction of developmental trajectories during zebrafish embryogenesis. Science 360, doi:10.1126/science.aar3131 (2018).

22. Coe, B. P. et al. Refining analyses of copy number variation identifies specific genes associated with developmental delay. Nat Genet 46, 1063–1071, doi:10.1038/ng.3092 (2014).

23. Kaplanis, J. et al. Integrating healthcare and research genetic data empowers the discovery of 49 novel developmental disorders. bioRxiv, 797787, doi:10.1101/797787 (2019).

24. Schizophrenia Working Group of the Psychiatric Genomics, C. Biological insights from 108 schizophrenia-associated genetic loci. Nature 511, 421–427, doi:10.1038/nature13595 (2014).

25. Stahl, E. A. et al. Genome-wide association study identifies 30 loci associated with bipolar disorder. Nat Genet 51, 793–803, doi:10.1038/s41588-019-0397-8 (2019).

26. Grove, J. et al. Identification of common genetic risk variants for autism spectrum disorder. Nat Genet 51, 431–444, doi:10.1038/s41588-019-0344-8 (2019).

27. Pardinas, A. F. et al. Common schizophrenia alleles are enriched in mutation-intolerant genes and in regions under strong background selection. Nat Genet 50, 381–389, doi:10.1038/s41588-018-0059-2 (2018).

28. Howard, D. M. et al. Genome-wide meta-analysis of depression identifies 102 independent variants and highlights the importance of the prefrontal brain regions. Nat Neurosci 22, 343–352, doi:10.1038/s41593-018-0326-7 (2019).

29. Finucane, H. K. et al. Partitioning heritability by functional annotation using genome-wide association summary statistics. Nat Genet 47, 1228–1235, doi:10.1038/ng.3404 (2015).

30. Finucane, H. K. et al. Heritability enrichment of specifically expressed genes identifies disease-relevant tissues and cell types. Nat Genet 50, 621–629, doi:10.1038/s41588-018-0081-4 (2018).

31. Langfelder, P. & Horvath, S. WGCNA: an R package for weighted correlation network analysis. BMC Bioinformatics 9, 559, doi:10.1186/1471-2105-9-559 (2008).

